# Micronutrient Optimization Using Design of Experiments Approach in Tissue Engineered Articular Cartilage for Production of Type II Collagen

**DOI:** 10.1101/2022.12.07.519522

**Authors:** Maria A. Cruz, Yamilet Gonzalez, Javier A. Vélez Toro, Makan Karimzadeh, Anthony Rubbo, Lauren Morris, Ramapaada Medam, Taylor Splawn, Marilyn Archer, Russell J. Fernandes, James E. Dennis, Thomas J. Kean

## Abstract

Tissue Engineering of cartilage has been hampered by the inability of engineered tissue to express native levels of type II collagen *in vitro*. Inadequate levels of type II collagen are, in part, due to a failure to recapitulate the physiological environment in culture. In this study, we engineered primary rabbit chondrocytes to express a secreted reporter, *Gaussia* Luciferase, driven by the type II collagen promoter, and applied a Design of Experiments approach to assess chondrogenic differentiation in micronutrient-supplemented medium. Using a Response Surface Model, 240 combinations of micronutrients absent in standard chondrogenic differentiation medium, were screened and assessed for type II collagen expression. Five conditions predicted to produce the greatest Luciferase expression were selected for further study. Validation of these conditions in 3D aggregates identified an optimal condition for type II collagen expression. Engineered cartilage grown in this condition, showed a 170% increase in type II collagen expression (Day 22 Luminescence) and in Young’s tensile modulus compared to engineered cartilage in basal media alone. Collagen cross-linking analysis confirmed formation of type II-type : II collagen and type II-type : IX collagen cross-linked heteropolymeric fibrils, characteristic of mature native cartilage. Combining a Design of Experiments approach and secreted reporter cells in 3D aggregate culture enabled a high-throughput platform that can be used to identify more optimal physiological culture parameters for chondrogenesis.

## INTRODUCTION

Osteoarthritis (OA) is the most common degenerative musculoskeletal disease and is projected to increase in prevalence^2,3^. OA is characterized by progressive degeneration of articular cartilage in the joints of the hands, knees, and hip due to an imbalance of cartilage anabolism and catabolism^2,3^. Articular cartilage is a form of specialized connective tissue, primarily composed of type II collagen, water, and proteoglycans with sparsely distributed chondrocytes^4^. Cartilage has limited healing and regenerative abilities given that its avascular nature limits access to circulating progenitor cells following physical insult^4,5^. Currently, there are no disease-modifying treatments for OA^2,5^. Available therapeutics offer short-lived relief of acute symptoms and do not prevent endpoint joint damage, therefore there is a strong need for the development of new disease-modifying therapeutics^2,5,6^.

Tissue engineering of cartilage has the potential to revolutionize the field by providing improved *in vitro* models for drug discovery and/or a biological replacement^6^. Tissue engineering incorporates the use of components such as cells, scaffolds, growth factors, and physical stimulation to generate biomimetic tissue^7^. However, tissue engineering of cartilage has been hampered by an inability to recapitulate the properties of native cartilage tissue, which we hypothesize is primarily due to insufficient type II collagen production. Whereas 90-95% of collagen in native tissue is type II collagen, several studies have reported much lower type II collagen levels in engineered tissue with values hovering around 20% despite modifications to increase collagen deposition^6,8-10^. We postulate that part of this deficiency in type II collagen is due to sub-optimal formulation of the culture medium used for cartilage engineering *in vitro*.

Differentiation media traditionally used for chondrocyte cell culture was noted to lack several micronutrients which are known to be physiologically essential for a host of biological processes^6^. Although the specific roles that some of these micronutrients have in chondrogenesis remain undefined, there are several findings that point towards these biomolecules having significant effects on cartilage generation and maintenance^6,11-16^. We hypothesize the addition of these vitamins and minerals to basal differentiation medium will promote type II collagen production *in vitro* and better mimic the physiological environment.

A Design of Experiments (DoE) approach was implemented in this study to screen different combinations of vitamins and minerals. DoE is a statistical technique that facilitates systematic optimization by producing experimental design models to study interactions of multiple factors on a desired outcome or response. DoE allows for a multi-factor, rather than a one-factor approach, that evaluates synergistic effects, and can predict optimal conditions while reducing the burden of conducting repetitive experiments. DoE has provided significant benefits to other fields of engineering and biotechnology but has rarely been used in cartilage tissue engineering and regenerative medicine^17,18^.

In this study, we have identified, for the first time, an optimal supplementation of physiologically necessary micronutrients to chondrogenic media, using a streamlined platform that includes a type II collagen promoter-driven *Gaussia* luciferase construct in primary rabbit articular chondrocytes combined with a DoE approach. This optimized chondrogenic media significantly enhances type II collagen expression in primary rabbit chondrocytes cultured in 3D cell aggregates and engineered cartilage sheets.

## RESULTS

### Stimulation of type II collagen by TGF-β1 in primary rabbit chondrocytes

To characterize the TGF-β1 response of engineered type II collagen promoter-driven *Gaussia* luciferase reporter (COL2A1-Gluc) in primary rabbit chondrocytes, cells were cultured in 3D aggregates in defined chondrogenic media supplemented with 0-10 ng/ml of TGF-β1, a known stimulator of type II collagen^6,19,20^. Conditioned media, containing the secreted *Gaussia* luciferase, was assayed for luminescence over three weeks. Dose response curves were generated from luminescence data at Day 7 (**Fig. 1a**) and Day 21 (**Fig. 1b**). As seen in Figure 1, there was a dose dependent increase in luminescence with a calculated 50% effective concentration (EC50) of 0.17ng/ml and 0.10ng/ml for Day 7 and Day 21 respectively.

**Figure 1:**
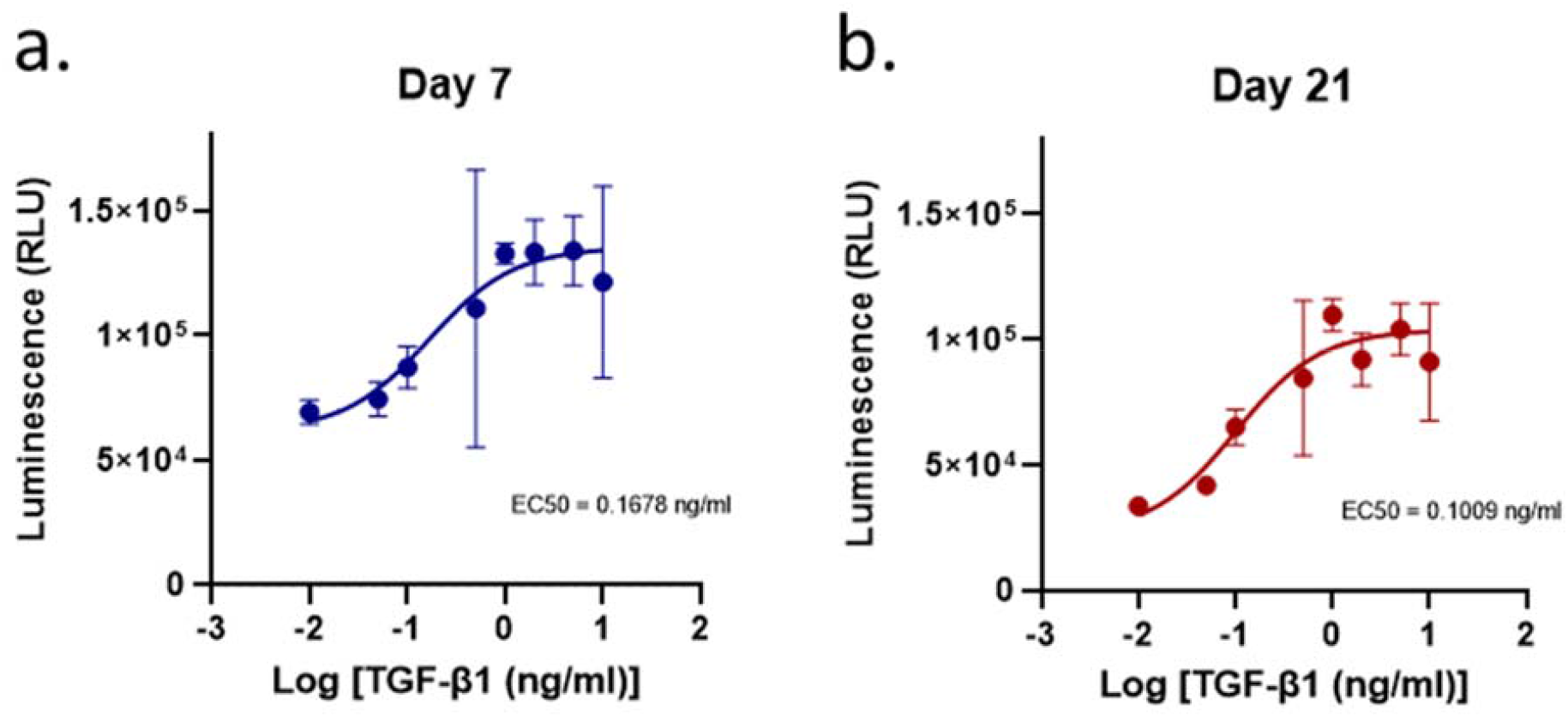
Dose effect of TGF-β1 on COL2A1-GLuc reporter rabbit chondrocytes. **a, b** Primary COL2A1-GLuc rabbit chondrocytes were grown in aggregate culture in the presence of different concentrations of TGF-β1 (0-10 ng/ml). Dose response curves were generated from transformed luminescence data at day 7 **(a)** and day 21 **(b)** after seeding. EC50s for each day are shown within each curve. Values are the mean ± S.D. n = 4.

### Response Surface Model and subsequent ANOVA analysis identified interactions between micronutrients that increased type II collagen promoter-driven expression of *Gaussia* luciferase

To identify potential interactions between factors and their effect on type II collagen expression, COL2-GLuc rabbit chondrocytes were seeded in 3D aggregate culture with DoE generated combinations of vitamins and minerals; media was sampled and replaced over three weeks. Combinations and concentrations are defined by the parameters set in the response surface model (**Table 1**) and are listed in **supplemental Table 2**.

**Table 1:**
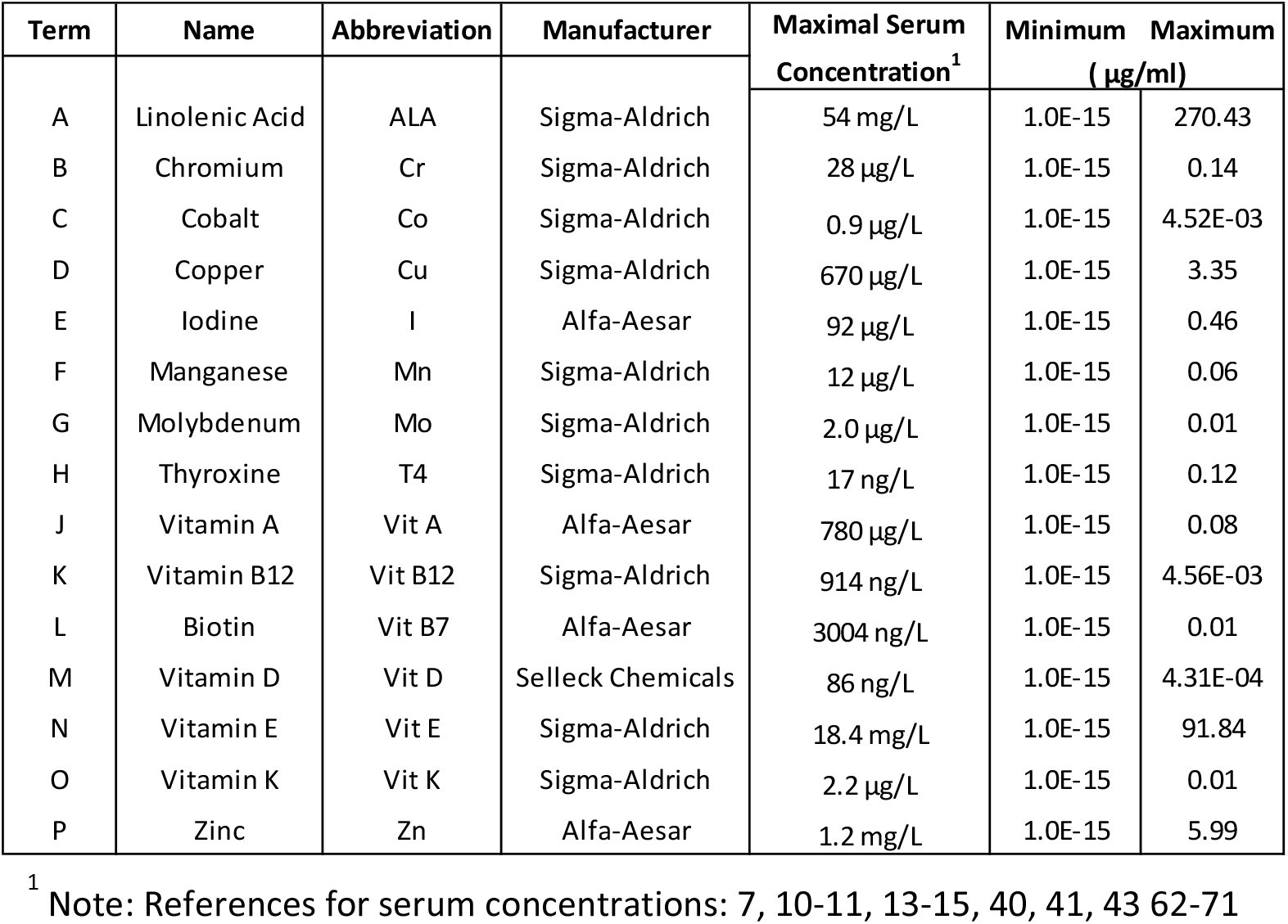
Concentrations of micronutrients absent in chondrogenic media with input parameters for DoE screen.

In the surface response model, each vitamin or mineral is introduced as an independent variable and is defined in Design-Expert (V.12, StatEase) as a model term. Luminescence signal over time, cumulative luminescence and resazurin data are defined as responses. The response surface study was designed as a quadratic model. **Fig. 2a-b** shows the normal probability plot after the data was transformed to fit the quadratic model for week 2 and week 3. The residuals are the deviation of each sample compared to its predicted value. For the residuals to be normally distributed they must show a linear trend, indicated by the red line, with little variation outside of it. As seen in **Fig. 2a-b** the residuals are normally distributed for all timepoints.

**Figure 2:**
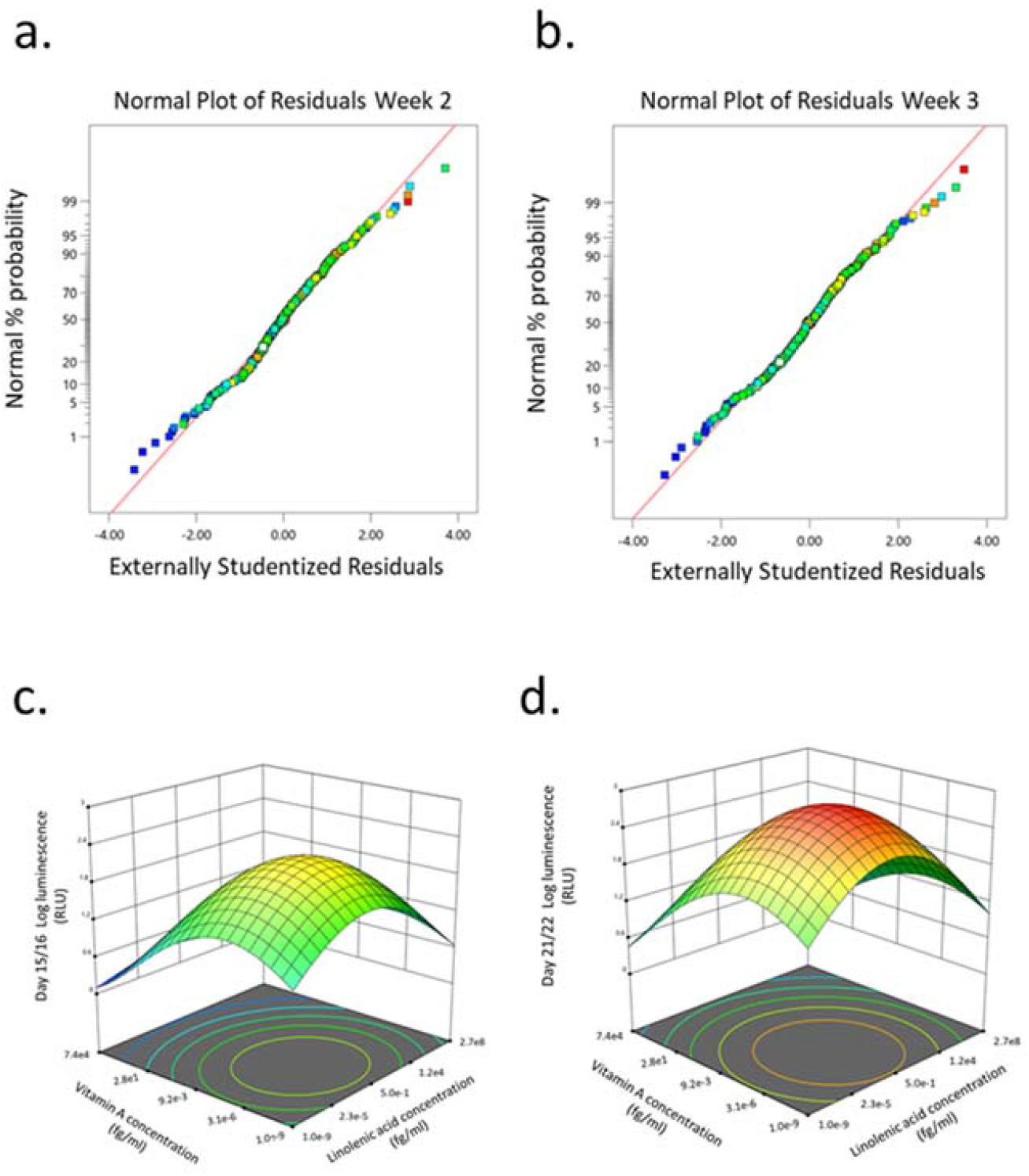
Dose effect of basal chondrogenic media supplemented with DoE micronutrient combinations on COL2A1-GLuc reporter Rabbit chondrocytes. Normal probability plots of the residuals for *Gaussia* Luciferase signal at weeks 2 (**a**) and 3 (**b**) after seeding. 3D surface response plots for interactive effects between vitamin A and linolenic acid at indicated weeks after seeding. End of week 2 **(c)**, and week 3**(d)**.

ANOVA analysis of luminescence expression for week 2 and week 3 of chondrogenesis identified significant model terms, i.e. factors that have significant effects on the responses (**Table 2**).

**Table 2:**
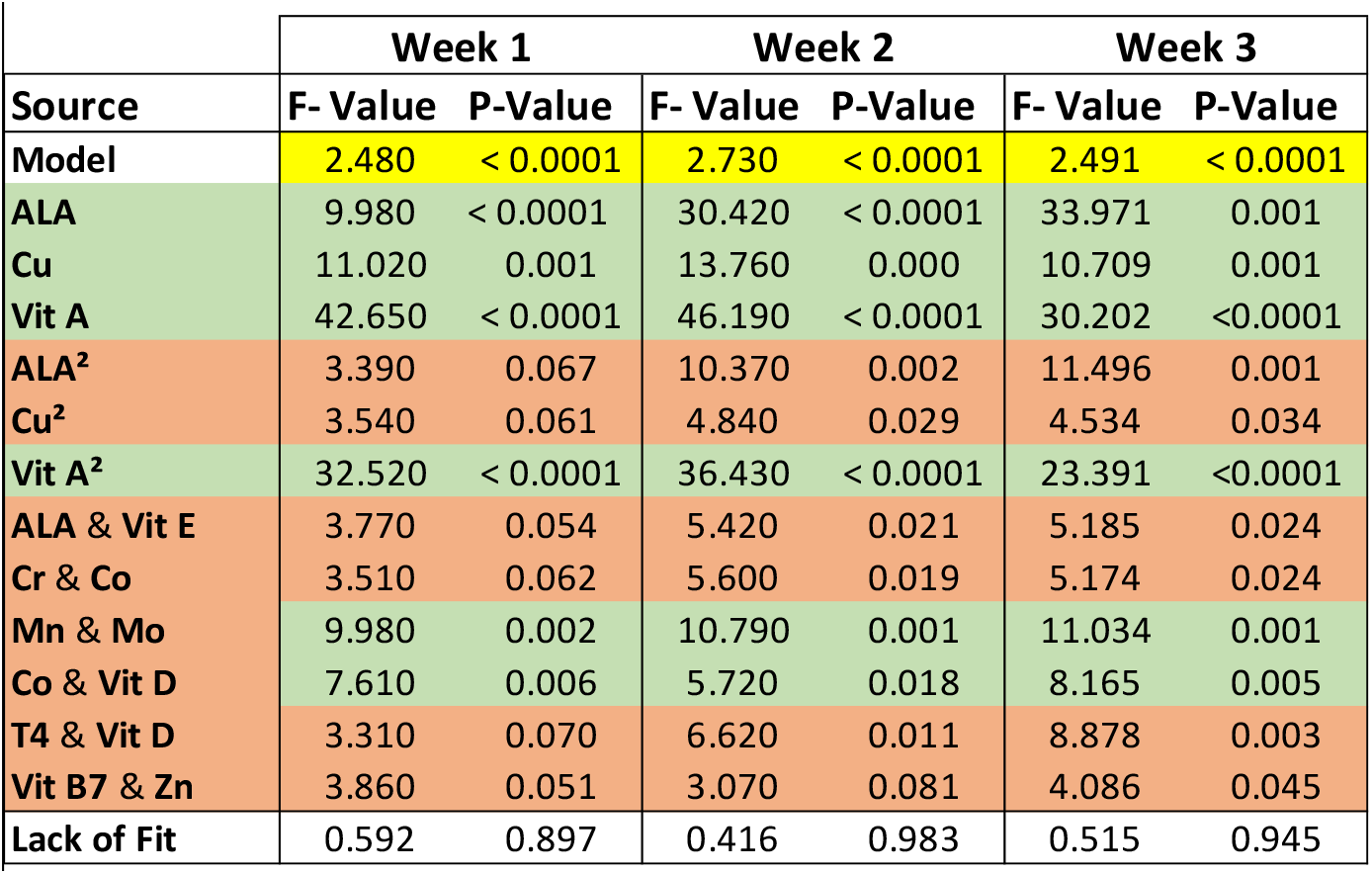
ANOVA analysis of surface response model to determine effects of micronutrients in COL2A1-GLuc reporter rabbit chondrocytes.

Results of this analysis include, F-values, P-values, and lack of fit test, which indicate how well the responses fit the model. As shown in **Table 2** the F- and P-values of the model, as well as the lack of fit test, over the three weeks support that the model is significant and thus the analysis for the associated terms is valid. **Table 2** displays model terms that were significant for at least one of the timepoints indicated shown by a P-value < 0.05. Significant single terms for all timepoints were linolenic acid, copper, and vitamin A. Several interactions were defined as significant for at least one of the timepoints shown (orange highlighted), while others including: chromium and cobalt, manganese and molybdenum, and cobalt and vitamin D were significant for all timepoints.

To contrast these results, testing factors in one factor at a time approach previously identified basal media supplemented with cobalt, chromium, thyroxine, or Vitamin B7, as having higher luminescence at multiple concentrations as compared to basal media alone (**Supplementary Fig. 1a**). Combinations of these factors, with factors previously shown to have an effect^6^, are shown in **Supplementary Fig. 1b**. Copper and Vitamin B7 alone or in combination with each other seemingly had no effect on luminescence, ∼40,000 RLU; however, in combination with thyroxine they significantly increased luminescence ∼70,000 RLU as compared to basal media alone, ∼40,000 RLU. This supports that there a synergistic effect between copper, biotin, and thyroxine. Furthermore, this supports the use of a Design of Experiments approach over a one factor a time approach as the DoE screen also identified factors that were significant in interactions but were not identified as individually significant.

Using a response surface model allows us to determine significant interactions between terms as well as determine and predict optimal concentrations of the terms within the parameters input into the initial model. 3D surface plots in **Fig. 2c-d** show the dose effect of two terms (linolenic acid and vitamin A) in relation to each other and to the response (luminescence) at week 2 (**Fig. 2c**) and week 3 (**Fig. 2d**) of chondrogenesis. During week 2 (**Fig. 2c**) there is a predicted optimal concentration for linolenic acid and vitamin A (approximately 3×10^−6^ fg/ml and 5 × 10^−1^ fg/ml respectively) that results in a maximal response that increases in week 3 (**Fig. 2d**), from 10^1.8^ RLU to 10^2.4^ RLU, although the optimal concentrations of the terms remain the same. Interestingly, the DoE model predicts that there is an optimal concentration for these in the femtomolar range (**Fig. 2c-d**) while each of the two factors, when analyzed as sole additives, were detrimental to chondrogenesis (**Supplementary Fig. 1a**).

**Fig. 2c-d** is a representation of only two of the terms and a predicted optimal for each individual timepoint. Through Design-Expert, multiple terms and responses can be analyzed together to derive predicted optimal concentrations for all terms. These predicted optima account for individual responses as well interactions between terms. Out of the generated predicted optimal conditions, 5 were selected for validation, designated as conditions 12, 25, 52, 72, and 89 (**Supplemental Table 2**).

### DoE predicted conditions improved *Gaussia* luciferase expression over basal media

To validate the Design-Expert generated predicted conditions, COL2A1-GLuc cells in aggregate culture were maintained in media supplemented with the predicted conditions (**Supplemental Table 2)** for three weeks. As seen in **Fig. 3a**, all conditions tested had increased luminescence over basal media control for all timepoints after day 8, which suggests that predicted conditions have an anabolic effect early in chondrogenesis. Cumulative luminescence seen in **Fig. 3b** is the sum of luminescence signal over all days in culture and confirms that increased luminescence at each timepoint results in an overall significant increase in type II collagen promoter-driven activity for all predicted conditions, with condition 25 having a higher cumulative signal of ∼1 × 10^6^ RLU as compared to basal media, ∼6 × 10^5^ RLU, and other predicted conditions. Single day luminescence shown for day 10 (**Fig. 3c**) and day 22 (**Fig. 3d**) supports an increase in luminescence that is statistically significant for all conditions tested as compared to basal media with condition 25 having an average luminescence signal that is twice of that in basal media, ∼2 × 10^5^ RLU vs 1 × 10^5^ RLU respectively, for both timepoints.

**Figure 3:**
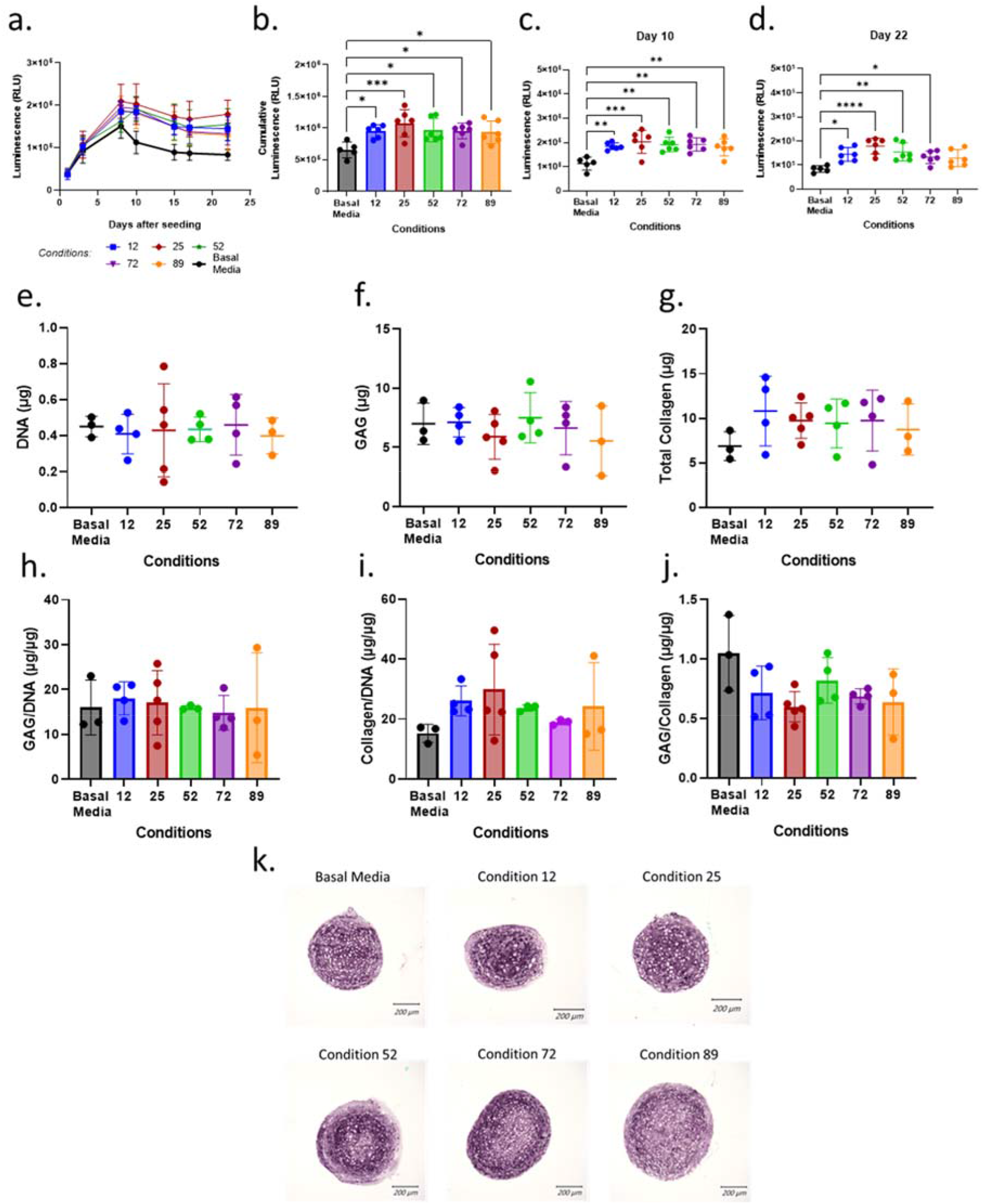
Validation of DoE predicted optimal conditions. **a-d** Conditions predicted by DoE analysis were tested in aggregate culture of COL2A1-GLuc reporter rabbit chondrocytes over 22 days. Results are shown as luminescence over 22 days sampled **(a)**, as well as cumulative luminescence signal **(b)**. To explore temporal effects, data was also analyzed at single day luminescence shown here for Day 10 **(c)** and Day 22 **(d)**. At day 22, aggregate cultures were assessed for total DNA **(e)**, glycosaminoglycan **(f)** and collagen **(g)** content. Results are also shown as total glycosaminoglycan and collagen normalized to DNA content **(h, i)** and to each other **(j)**. Alternatively, aggregates were fixed, embedded in paraffin and sectioned. **k** Sections were analyzed for type II collagen. Scale Bars, 200um. **a-d** N = 6. **e-j** N = 5. Replicates or means ± S.D. and ** p <0.01 and *** p <0.001 vs. Basal Media control.

To corroborate results seen by luminescence output, endpoint biochemical assays were performed at day 22 of the experiment to quantitate DNA, glycosaminoglycan (GAG) and total collagen content. **Fig. 3e** shows an average of ∼0.4 μg of DNA per sample with no significant difference between conditions tested, which suggests that predicted conditions have no effect on cell proliferation or viability over 22 days. Total glycosaminoglycan content is shown in **Fig. 3f** and as amount per microgram of DNA in **Fig. 3h**. As expected, DoE predicted conditions did not significantly affect glycosaminoglycan production over 22 days, although condition 89 shows large variability between samples as compared to other conditions. Quantification of total collagen, as seen in **Fig. 3g** shows an increase in aggregates cultured in predicted conditions to ∼10 μg over basal media (∼6 μg). However, when normalized to micrograms of DNA (**Fig. 3i**) only conditions 12, 25 and 89, show increased collagen as compared to aggregates cultured in basal media with significant variability within each group. Glycosaminoglycan to collagen ratio (**Fig. 3j**) further support an increase in collagen with no change in GAG content. Immunohistochemistry for type II collagen (**Fig. 3k**) confirms the presence of type II collagen for cell aggregates in all conditions at day 22 with similar staining pattern across all conditions.

### No single factor from optimized condition 25 significantly impacts type II collagen stimulation in primary rabbit chondrocytes

When using a one factor at a time approach, thyroxine (T4) was required in the tested combinations for type II collagen stimulation over basal media (**Supplementary Fig 1b**). To test if one factor was solely responsible for the enhancement in luminescence seen over basal media, aggregates were cultured in the predicted DoE condition 25, or combinations where one factor was removed from condition 25 for 22 days. **Fig. 4a** represents luminescence data for all days tested with basal media shown by a thicker black line and complete condition 25 shown by the thicker red line. All conditions had increased luminescence as compared to basal media after day 8. For statistical analysis, day 10 luminescence (**Fig. 4b**) is shown for all conditions tested.

**Figure 4:**
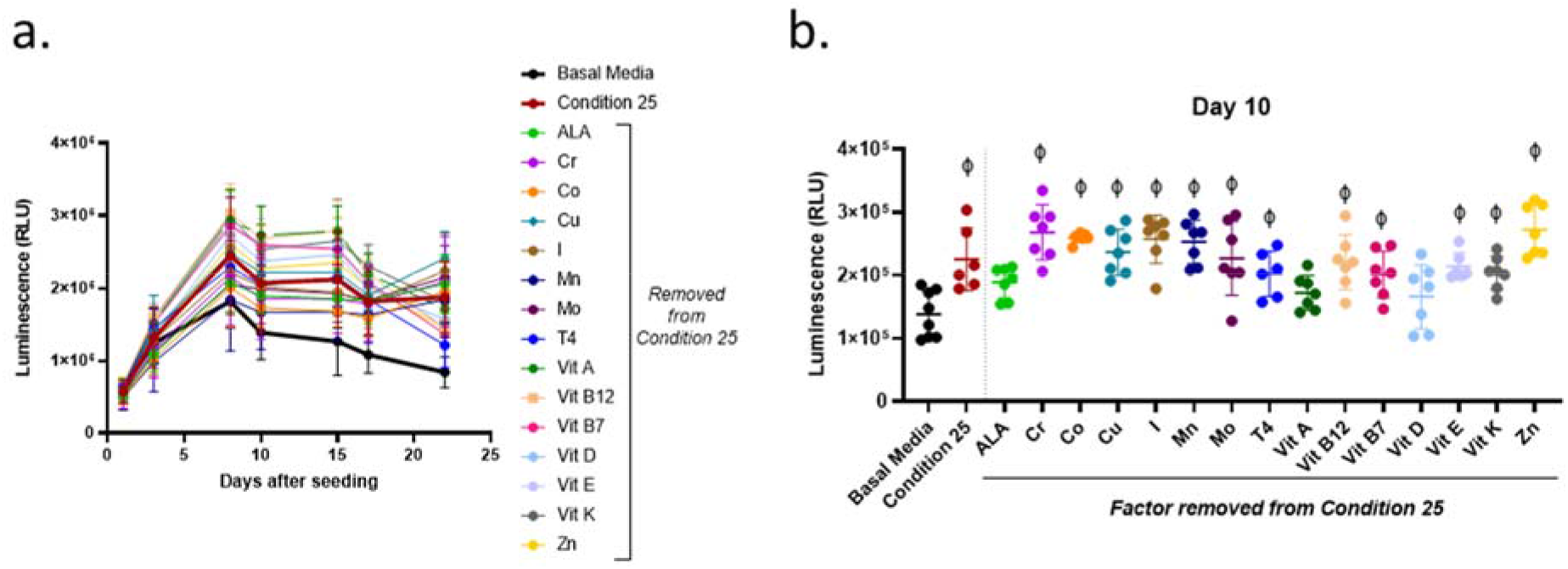
Effect of a single micronutrient removal from DoE predicted condition 25 on type II collagen driven expression of *Gaussia* Luciferase. Aggregates of COL2A1-GLuc primary rabbit chondrocytes were cultured with condition 25 or condition 25 with a single micronutrient removed as indicated. Results are shown as luminescence over 22 days **(a)**. Luminescence is shown for a single timepoint, Day 10 **(b)**. ϕ indicates p <0.05 vs. Basal Media control. N = 7. Mean ± S.D.

Aggregates cultured in all conditions had increased luminescence, however, effects by condition 25 with Linolenic acid, vitamin A or with vitamin D removed, were not statistically significant as compared to basal media, suggestive of a major role for these biomolecules. Formulations where any of the other factors were removed from Condition 25 had significant increases in luminescence as compared to basal media alone. There was no statistical significance between any of the conditions with a single factor removed as compared to complete condition 25. These results are evidence that no single factor within DoE predicted condition 25 is solely responsible for the higher stimulation of type II collagen promoter activity observed.

### Relative concentrations of vitamins and minerals, regardless of absolute concentrations, in DoE predicted conditions play a significant role in type II collagen stimulation in primary rabbit chondrocytes

To determine if ratios, regardless of the absolute concentration, within the DoE predicted conditions were sufficient for type II collagen stimulation, aggregate cultures were treated with the DoE predicted conditions at the concentrations given or at 1/15^th^ of the predicted concentrations. Statistical analysis of luminescence at Day 10 (**Fig. 5a**) and Day 17 (**Fig. 5b**) showed no significant differences between the conditions at 1x or 1/15x of their DoE predicted concentrations.

**Figure 5:**
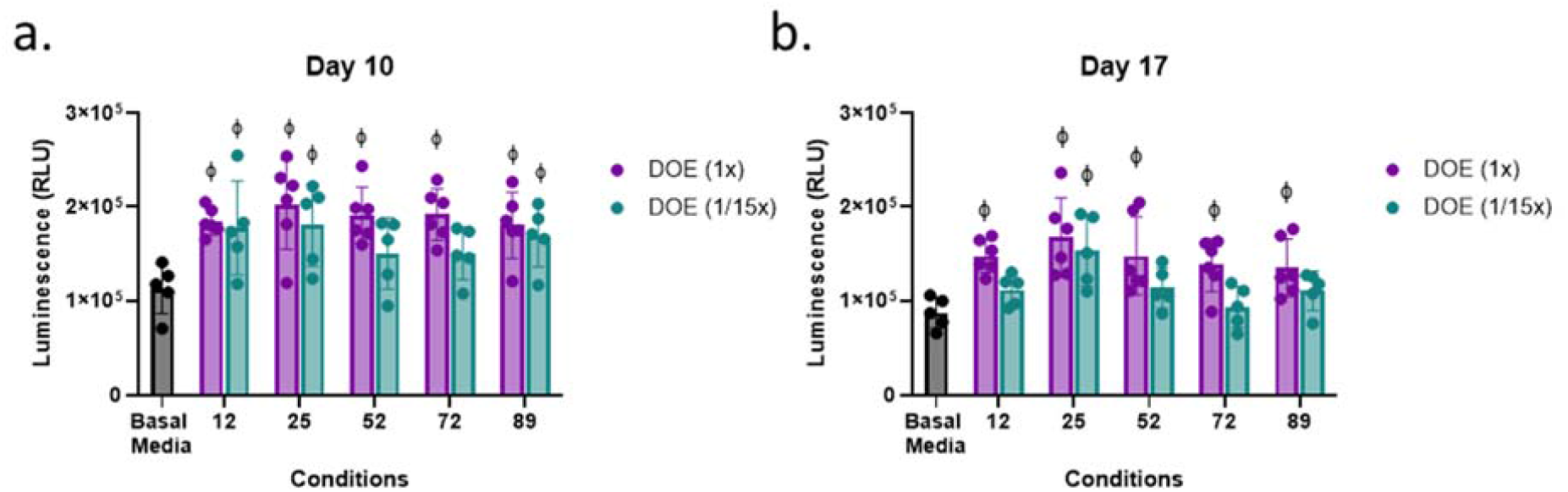
Effect of absolute versus relative concentration of micronutrients on Type II collagen driven expression of *Gaussia* Luciferase. Aggregates of COL2A1-GLuc primary rabbit chondrocytes were cultured with predicted DoE conditions (1x) or the same ratios of these conditions at 1/15th the optimal predicted concentration (1/15x) over 22 days. Luminescence results are shown for day 10 **(a)** and day 17 **(b)**. ϕ indicates p <0.05 vs. Basal Media control. N = 5 for DoE at 1/15x and N = 6 for DoE at 1x. Individual replicates and the mean ± S.D. are shown.

Interestingly, while aggregates cultured at the DoE predicted conditions had significantly higher luminescence as compared to those in basal media, aggregates cultured in most conditions at 1/15^th^ did not. Only condition 25 at 1/15^th^ the concentration showed a significantly higher level of luminescence at both days vs. Basal Media, ∼1.5 to ∼1.7 ×10^5^ RLU vs ∼1 × 10^5^ RLU. This suggests that the combinatorial effect of the vitamins and minerals plays a significant role in type II collagen stimulation in primary rabbit chondrocytes, but concentrations as predicted by the DoE are optimal for type II collagen stimulation.

### DoE predicted condition 25 stimulates type II collagen in tissue engineered rabbit cartilage

To determine if condition 25, the best performing DoE predicted condition, could have an effect in cartilage tissue engineering, we cultured COL2A1-GLuc or Non-transduced (NonTr) primary rabbit chondrocytes in custom, 3D printed bioreactors adapted from Whitney GA, et al. shown in **Fig 6a-b** over 22 days^21,22^.

**Figure 6:**
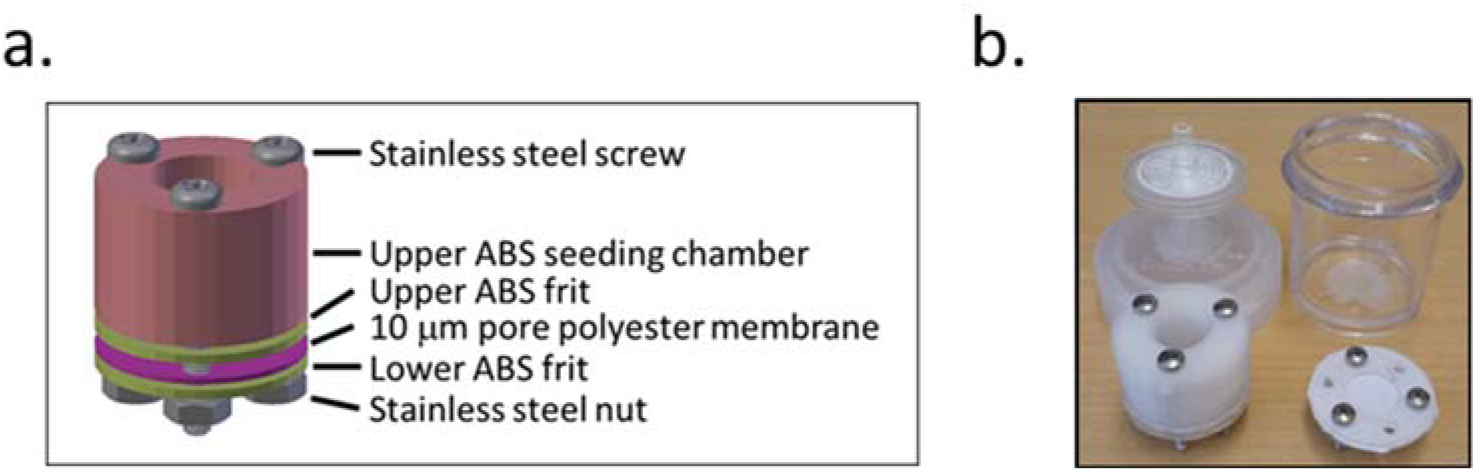
Biochamber for the generation of tissue engineered cartilage sheets Custom 3D printed ABS biochambers were designed to generate tissue engineered articular cartilage sheets. **a** Photo of printed biochamber and Nalgene container fitted with sterile filter top. **b** Model of the biochamber assembly.

At the end of this culture period, 1.2cm^2^ cartilage sheets were collected as shown in **Fig. 7a**. Over the 22 days bioreactors housing COL2A1-Gluc cells were regularly sampled for luminescence. **Fig. 7b** represents luminescence data for all days tested.

**Figure 7:**
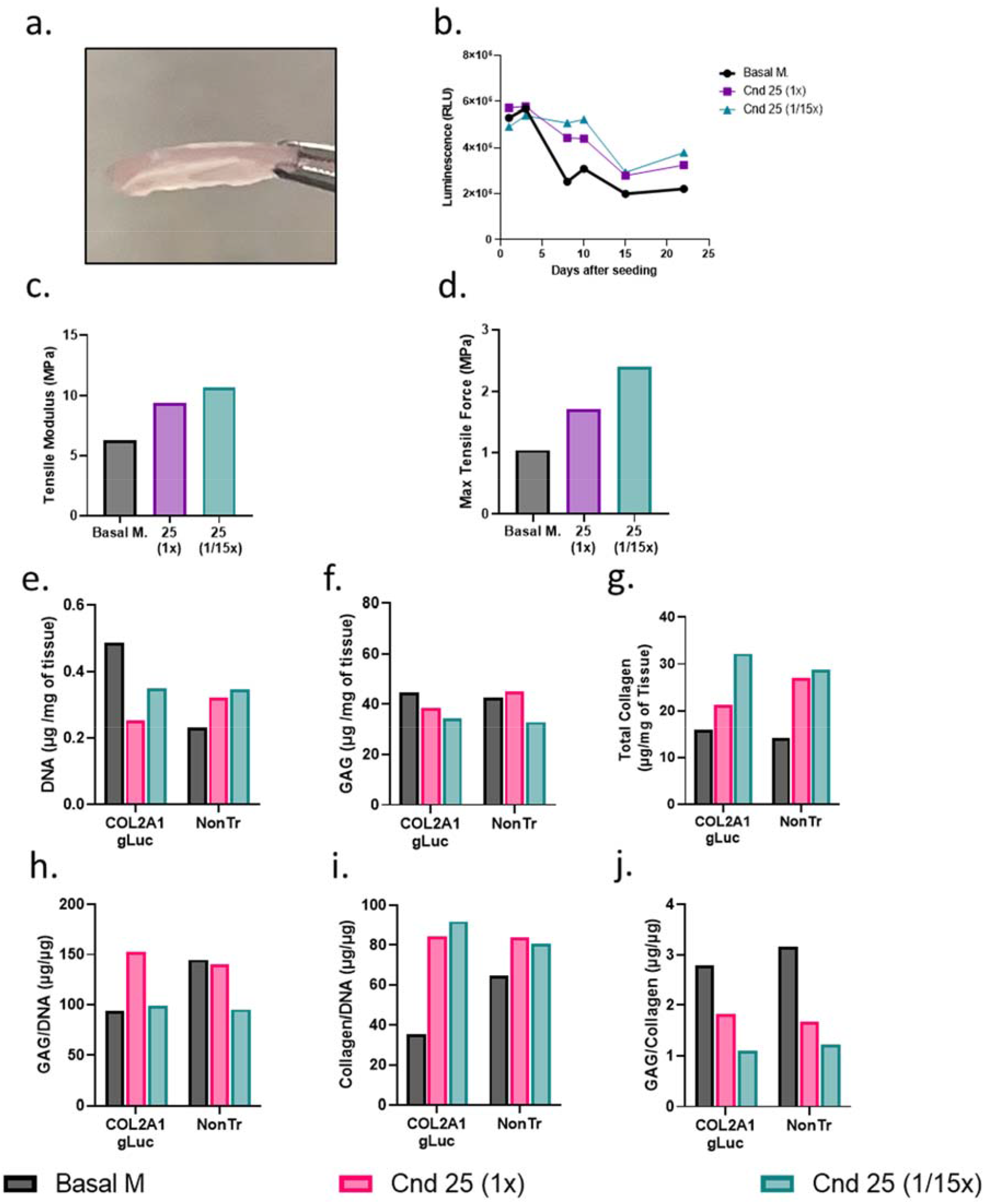
Tissue engineered cartilage sheet response to supplementation with condition 25. COL2A1-GLuc primary rabbit chondrocytes were cultured in bioreactors with condition 25 (1x) or condition 25 at 1/15th the concentration (1/15x) over 22 days. **a** On day 22, engineered sheets were collected. **b** Luminescence signal over 22 days in culture. Max tensile force **(c)** and tensile modulus **(d)** are shown. DNA **(e)**, glycosaminoglycan **(f)** and collagen content **(g)** of the tissue were assessed. Data is also calculated as GAG/DNA **(h)**, collagen/DNA **(i)** and GAG/collagen **(j)**.

Similar to results seen when conditions were tested in aggregates (**Fig. 3**), both condition 25 at 1x and at 1/15^th^ concentration showed increased luminescence as compared to basal media after day 8. A different curve in luminescence is observed in **Fig. 7b** as compared to **Fig. 3a** because samples were collected from inside of the biochamber for day 1 and day 3. After day 3, medium was added to increase exchange between the inside and outside of the chamber, thus samples collected are from a larger volume resulting in a decrease in luminescence from day 3 to day 8. After sheets were collected, several biopsy punches of the sheets were obtained for endpoint analysis. To determine whether increased COL2A1 reporter activity translates to improved mechanical properties of engineered cartilage, we assessed tissue elasticity via tensile testing. As expected, we observed a marked increase in the Tensile modulus (**Fig. 7c**) as well as maximal tensile force (**Fig. 7d**), in sheets generated in condition 25 media at 1x and 1/15^th^ of the concentration as compared to basal media, from ∼6 MPa in basal media to ∼8 MPa in condition 25 and ∼10 MPa in condition 25 at 1/15^th^ concentration. In contrast, compression testing resulted in a marked decrease in stiffness for sheets generated in conditions 25 regardless of concentration (**Supplementary Fig. 2**).

To evaluate whether the luminescence results reflected matrix accumulation, biochemical assays were performed at day 22 of the experiment to quantitate DNA, glycosaminoglycan (GAG) and total collagen content from bioreactors seeded with COL2A1-Gluc and non-transduced cells. Both COL2A1-gLuc and non-transduced cell generated sheets had similar trends in DNA (**Fig. 7e**), GAG (**Fig. 7f**), and total collagen (**Fig. 7g**) except for the sheet generated with COL2A1-Gluc cells in condition 25 at (1x), which shows substantially lower collagen as compared to its non-transduced counterpart (**Fig. 7g**). Sheets in supplemented groups had overall higher collagen content per milligram of tissue, as well as per microgram of DNA (**Fig. 7g** and **7i**). Total glycosaminoglycan content is relatively constant for all groups at ∼40 μg per mg of wet tissue weight (**Fig. 7f**), when normalized to collagen content (**Fig. 7j)** there is a noticeable decrease in sheets generated in condition 25 (both 1x and 1/15x) suggesting that condition 25 specifically affects collagen content and not glycosaminoglycan content. Immunofluorescence of sections of the engineered cartilage confirmed the presence of type II collagen (**Fig 8a**). Interestingly, condition 25 (1/15^th^) had a more extensive distribution of type II collagen signal as compared to basal media alone which showed an uneven pattern of staining. Collagen heteropolymer analysis was also carried out on samples of the engineered cartilage sheets and compared to native rabbit articular cartilage to further analyze collagen content. **Figure 8b** shows that the major pepsin-resistant Coomassie blue-stained band in both transfected and non-transfected articular chondrocyte cultures migrated identically to the α1(II) chain of type II collagen in the adult rabbit articular cartilage. β1(II) chains (dimers of α1(II) chains) were also observed in all lanes (**Fig. 8b)**. Two faintly stained bands migrating slightly slower than that of the α1(II) chains, are the α1(XI) and α2(XI) chains of type XI collagen as previously identified by mass spectrometry and described by McAlinden et al.^23,24^. The α1(I) and α2(I) chains of type I collagen were neither detected in engineered cartilages nor in the control articular cartilage (if present the α2(I) chain migrates slightly faster than the α1(II) chain), indicating that type II collagen and type XI collagen were the major collagens synthesized by the cultured chondrocytes and the cartilage collagen phenotype was maintained **(Fig. 8b)**. The western blot shown in **Fig. 8c** confirms that the Coomassie blue stained bands were indeed α1(II) chains of type II collagen.

**Figure 8:**
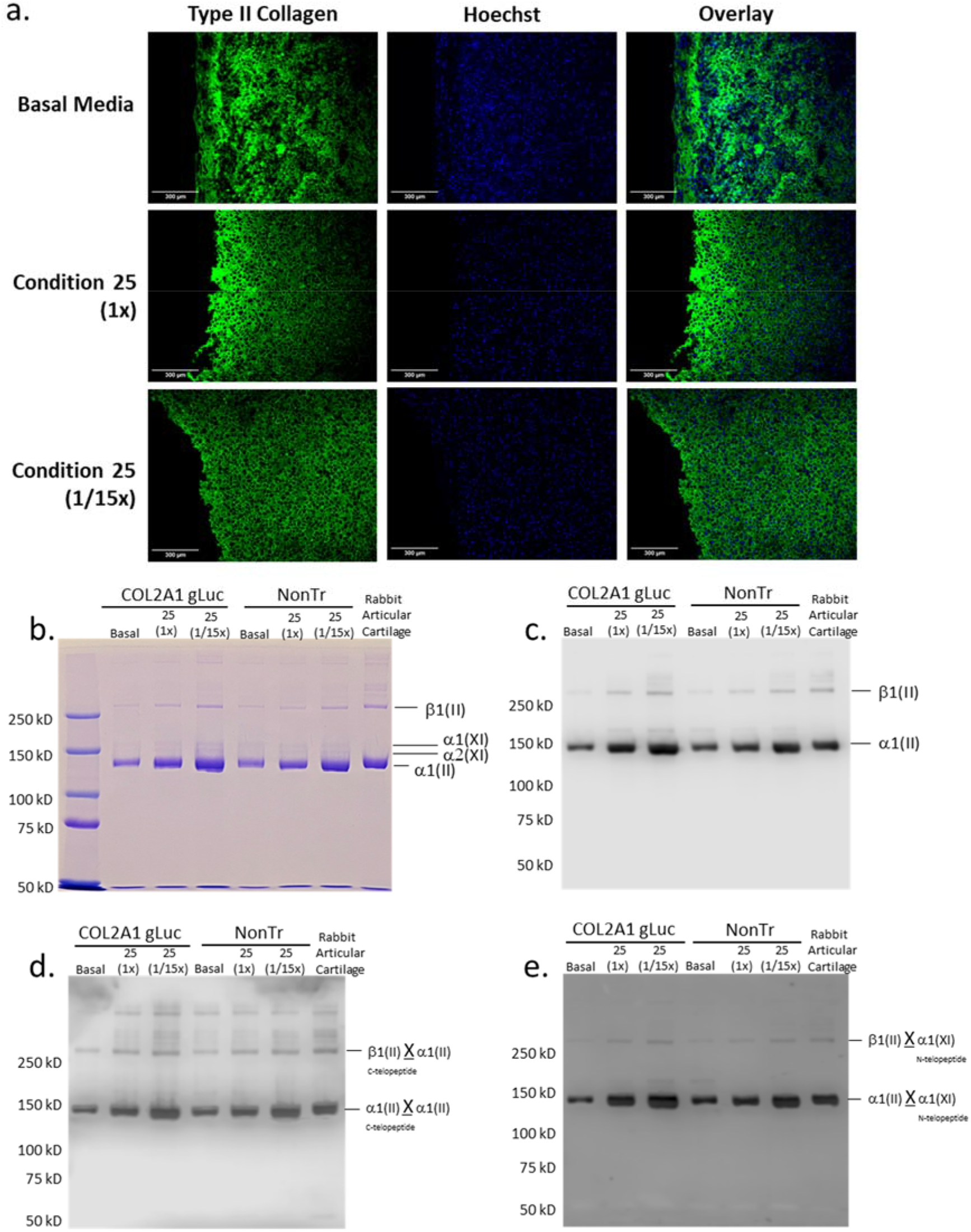
Analysis of Type II collagen and heteropolymer formation with Type IX collagen in tissue engineered cartilage. **a** Engineered cartilage was analyzed for type II collagen. Scale Bars, 300um. **b** Coomassie blue-stained SDS-PAGE gel of pepsin solubilized collagen showing β1(II), α1(XI), α2(XI) and α1(II) chains. Equivalent dry weight (25 μg) was loaded. **c** Western blot of samples equivalent to those in **(b)** and probed with anti-type II collagen antibody (1C10). **d** Western blot of samples equivalent to those electrophoresed in **(b)** (above) and probed with mAb 10F2. This antibody specifically recognizes the C-telopeptide domain of type II collagen when it is cross-linked to another α1(II) collagen chain. **e** Western blot of samples identical to those in **(b)** probed with antibody 5890. This antibody specifically recognizes N-telopeptide domain of α1(XI) collagen when cross-linked to chains of α1(II) and β1(II). **X** denotes crosslinks.

Using mAb 1C10, which specifically recognizes type II collagen chains^25^, intense staining of both the α1(II) and β1(II) chains were observed in all the cultures and the adult rabbit cartilage. Since equivalent engineered cartilage dry weights loads were electrophoresed in all the lanes, densitometry of the Western blot revealed increased levels of type II collagen (α1(II) + β1(II)) reactivity under condition 25 in both transfected and non-transfected cultures compared to basal conditions. Condition 25 (1/15x) however showed highest levels of reactivity indicating highest type II collagen retained in the Condition 25 (1/15x) sheet. This is consistent with the analytical results in **Fig. 7g** showing highest collagen content under this condition on a per mass basis. Using a refined Western blot method^21^ we were able to identify precise domains of collagen chains that were cross-linked in these cultures. As seen in **Fig. 8d**, western blotting using mAb 10F2^24,26^ recognized the α1(II) and β1(II) chains in all the cultures and the adult rabbit cartilage. This is evidence that the C-telopeptide of the α1(II) chain specifically recognized by this antibody was cross-linked to the helical lysine (K87) residues in a fraction of α1(II) collagen chains and, thus, type II to type II collagen cross-links had formed in these cultures^21^. It must be reiterated that pepsin-extracted α1(II) collagen chains are devoid of telopeptides unless they are cross-linked to the lysine residues in the helical regions of α1(II) chains^21,24^.

To examine if type II and type XI collagen molecules in these cultures were stabilized by these cross-links, we used the pAb 5890^23,24^. As seen in **Fig. 8e**, this antibody also recognized the α1(II) chains and the β1(II) chains of type II collagen in the tissue engineered sheets and adult rabbit cartilage. As shown before^24^, this means that the N-telopeptide of the α1(XI) chain to which this antibody was raised was cross-linked to the helical lysine (K930) residue in a fraction of α1(II) chains of type II collagen molecules and thus a hetero-polymer of type II and type XI collagens had formed in all these cultures. A faint reactivity of the α1(XI) chain was observed in some of the engineered cartilage cultures that probably indicates that N-telopeptides of α1(XI) chain are cross-linked to helical lysine of another α1(XI) chain and a homo-polymer in a fraction of type XI collagen had also formed in these cultures. The data confirms that a polymer of type II collagen had formed in tissue engineered cartilage sheets and a mature collagenous heteropolymer of cross-linked type II-XI collagen fibrils had formed.

## DISCUSSION

Our previous efforts to optimize media conditions tested the effect of 15 different micronutrients and thyroxine on murine chondrocytes using proposed concentrations based on physiologic levels in a one-factor at a time approach^6^. In that work, we identified copper, vitamin A and linolenic acid as having a positive effect on chondrogenesis. Overall, we found that combinations of these micronutrients were able to increase the expression of type II collagen when tested temporally and in a dose-dependent manner^6^. While we showed that vitamins and minerals affect type II collagen production in murine chondrocytes *in vitro*, we noted several limitations of using a traditional one factor at a time approach. This approach consists of experimental runs that are executed to hold every factor constant except for the variable of interest. This approach poorly reflects the complexity of *in vivo* conditions by failing to account for important interactions and largely relies on iterative experiments and trial and error for optimization. In the current study, we combined a non-destructive reporter, primary rabbit chondrocytes, 3D culture in 96-well plates, automated pipetting, and Design of Experiments approach as an efficient high throughput platform. We were able to not only identify interactions of micronutrients that had an effect on type II collagen expression, but to also derive an optimal combination containing all missing factors as a supplement to traditional basal media, that simultaneously increased type II collagen expression.

There are several advantages to the platform we implemented in this study. 1) we made use of primary rabbit articular chondrocytes as a model for healthy cartilage. Primary cells have the advantage of being more relevant in orthopedic research than cell lines and thus more likely to mimic responses *in vivo*^27-30^. 2) using 3D cell aggregates adapted to 96-well plates cultured in physioxic conditions, which we adapted for use in an automated system, allows chondrocytes to maintain their phenotype as compared to 2D culture^31-35^. 3) We made use of a secreted *Gaussia* Luciferase reporter^36-40^. Traditional biochemical assays to evaluate chondrogenesis typically rely on destructive endpoint analysis, and due to the length of culture of the samples, low cell number in aggregates, and long and laborious processes, can result in high variability as seen in **Fig. 3e-j**. Using a secreted reporter allows us to sample the media without lysis of the aggregate and thus provides the ability to examine the temporal effects of the treatment conditions on chondrogenesis. Because media is replaced every 2-3 days *Gaussia* luciferase readings provide a readout of the activity of the type II collagen promoter at early, mid and late timepoints in chondrogenesis. This was seen and confirmed by the TGF-β1 dose response curve where we have shown, for the first time, that the effective dose of TGF-β1 on type II collagen stimulation differs throughout the process of chondrogenesis. Furthermore, type II collagen expression levels are 66% higher at early timepoints with a decrease in activity at later timepoints of chondrogenesis (**Fig. 1a** and **1b**). The *Gaussia* luciferase assay is simple, sensitive and fast to perform and thus reduces variability between samples.

Using this platform, we successfully screened 240 combinations of vitamins and minerals for their ability to promote type II collagen. Previous studies have shown that several of the micronutrients we tested can play a role in chondrogenesis^6,8,13,41-44^. Previously, vitamins D and K were shown to play a role in the development and regulation of chondrogenesis, while vitamin A exhibited inhibitory action on *in vitro* chondrogenic differentiation^12,42^. Additionally, vitamin E has exhibited oxidative stress inhibition during *in vivo* and clinical studies^13^. Other trace minerals such as copper and zinc promote extracellular matrix formation and deficiencies in selenium and iodine have been shown to impair bone and growth formation^44^. Molecules like linoleic acid are known to enhance the metabolic activity of differentiating cells, while thyroxine was shown to increase type II collagen expression and glycosaminoglycan (GAG) deposition in scaffold-free engineered cartilage tissue^8,12^. In this study we identified linolenic acid, copper, and vitamin A, as well as interactions between various vitamins and minerals **(Table 2)** as having significant effects on type II collagen stimulation.

These findings relied on the use of Response Surface Methodology based on Design of Experiments which significantly reduces the number of trials, accounts for errors in the model, and for interactions between factors^45-48^. Statistical analysis with this approach allowed us to predict an optimal combination of vitamins and minerals, condition 25, that when tested *in vitro* showed significant increases in type II collagen as compared to the basal media control. Furthermore, we removed one factor at a time from condition 25 and confirmed the importance of linolenic acid, vitamin A and vitamin D and their interactions in type II collagen stimulation, further confirming the validity of the DoE results.

While Response Surface Methodology has significant advantages, its effectiveness does rely on the data fitting a second order polynomial model, thus fit statistics are crucial to ensure that the data fits the model^47,49^. In addition, the validation of any findings is essential. Other aspects to consider include parameter selection for optimization of factors and response, as well as examining predicted values before validation. In our study, this is seen by our predicted condition 25, while having a predicted desirability of 0.595, it was selected for validation due to the high predicted individual responses during analysis. When tested *in vitro*, it showed similar if not better responses than other selected conditions. There are few studies that have used Design of Experiments to look at biological processes, typically investigating fewer factors^17,18,50^. To date, this study is the first to apply a response surface model to primary chondrocytes.

After validation of condition 25 in aggregates we explored this supplementation in tissue engineered cartilage. We used custom 3D printed bioreactors adapted from Whitney, GA et al. to generate cartilage sheets *in vitro* (**Fig. 6** and **Fig. 7a**)^21,22,51^. Similar to our findings in cell aggregates, type II collagen promoter-driven expression of *Gaussia* Luciferase was significantly increased as compared to cells in basal media in engineered cartilage. Biochemical studies supported an increase in total collagen content. Western blots of pepsin extracted samples confirm the increase is type II collagen, specifically, in sheets supplemented with condition 25 (**Fig 8c**). Collagen x-link analysis supports the formation of type II collagen to type II collagen and type II collagen to type IX collagen heteropolymers, as in native rabbit cartilage (**Fig 8d** and **Fig 8e**). These crosslinks are characteristic of mature cartilage. This is significant as cell processes, particularly in tissue engineering, are often context dependent^52-54^. It is interesting to note that condition 25 at 1/15^th^ was optimal for type II collagen expression as compared to condition 25 at 1x, as seen by luminescence, immunofluorescence and western blot. It is possible that higher concentrations are not needed by chondrocytes and could even be detrimental for chondrogenesis resulting in greater type II collagen expression when the concentrations are decreased. Multiple cell types, like osteoblasts, endothelial cells and vascular smooth muscle cells, have specialized mechanisms to recycle and fully utilize vitamins and minerals, as these cannot be synthesized by humans^55-58^. Investigation of micronutrient recycling in chondrocytes has not been well studied and was beyond the scope of this work.

Supplementation with condition 25 also altered the mechanical properties of the engineered cartilage. While it increased the tensile modulus of engineered cartilage, unexpectedly, we observed a decrease in Young’s modulus in compressive tests as compared to basal media (**Supplementary Fig. 2**), suggestive of decreased stiffness. While it is thought that type II collagen generally increases the tensile properties of cartilage, there is no clear correlation between type II collagen and stiffness^59^. Furthermore, mechanical testing of live biological tissue is also confounded by the method of testing. Patel JM et al.^60^, has explored the inconsistencies present with various modes of mechanical testing which make any comparison of our findings to previous literature extremely difficult. Despite a decrease in the compressive modulus the engineered cartilage generated with condition 25 shows mechanical and biochemical properties closer to that of native cartilage than engineered cartilage generated in basal media alone.

## CONCLUSIONS

This study demonstrates that the physiologic environment of micronutrients to culture chondrocytes has a far greater impact on chondrogenesis than previously appreciated.

Supplementation of culture medium with 15 micronutrients, that are physiologically present in the articular joint, can be tailored to improve *in vitro* chondrogenesis, and the biochemical and mechanical properties of tissue engineered cartilage. Our results show that the presence and concentrations of seemingly minor components of culture medium can have a major impact on chondrogenesis. Furthermore, we established a streamlined process using Design of Experiments and primary reporter chondrocytes as a way to identify optimal chondrogenic conditions *in vitro*.

## METHODS

### Rabbit Primary Chondrocyte Isolation

Rabbits were euthanized under American Veterinary Medical Association guidelines and knees were isolated within 2 hours of euthanasia. The articular knee joints were dissected under sterile conditions, and articular cartilage was isolated from both the femoral condyle and the tibial plateau. Isolated cartilage was diced into <1mm^3^ pieces before sequential digest, first in hyaluronidase for 30 min (660 Units/ml Sigma, H3506; in DMEM/F12 with pen/strep/amphotericin B, 30ml), followed by collagenase type II for ∼16 hours at 37°C (583 Units/ml Worthington Biochemical Corp.; in DMEM/F12 with 10% FBS, 1% pen/strep/fungizone, 30ml). The digest was then filtered through a 70 μm cell strainer, washed with DMEM/F12, and resuspended in growth media (DMEM/F12 supplemented with 10% FBS, 1% pen/strep). Cells were subsequently infected as described below or cryopreserved (95% FBS, 5% DMSO).

### Lentiviral Construct

Lentivirus was generated as previously described^6^. Briefly, an HIV based lentiviral third generation system from GeneCopoeia was used to generate pseudolentiviral particles. Custom ordered COL2A1-*Gaussia* Luciferase plasmid (HPRM22364-LvPG02, GeneCopoeia, Inc.), envelope (pMD2.G) and packaging (psPAX2) plasmids were amplified in *Escherichia coli* (GCI-L3, GeneCopoeia) and silica column purified (Qiagen Maxiprep) before being co-transfected into HEK293Ta (GeneCopoeia) cells via calcium chloride precipitation. Pseudolentiviral particles were harvested from conditioned media after 48h and concentrated via ultracentrifugation (10,000 RCF, 4°C, overnight). Titers for COL2A1-Gluc lentivirus were estimated via real-time PCR and aliquots stored at -80°C.

### Lentivirus Infection of Primary Rabbit Chondrocytes

Isolated rabbit chondrocytes were seeded at 6,200 cells/cm^2^ in growth media and allowed to adhere overnight (∼20% confluency). Cells were infected with lentivirus (COL2A1-GLuc; MOI 25 in growth media) in the presence of 4μg/ml polybrene (Opti-mem, Gibco) for 12h. Lentiviral medium was replaced with growth medium and cells expanded to ∼90% confluency. Cells were subsequently plated on flasks coated in porcine synoviocyte matrix^5,61^ and selected with puromycin (2 μg/ml) when 70% confluent for 48 hours. Culturing of rabbit chondrocytes during infection was done in physioxic (37°C, 5% O_2_, 5% CO_2_) conditions. Newly generated COL2A1-GLuc cells were cryopreserved at the end of this first passage (95% FBS, 5% DMSO). These cells were used for all subsequent studies.

### Chondrogenic Culture

Rabbit COL2A1-GLuc were thawed and seeded in growth media at 6000 cells/cm^2^ and expanded to 90-100% confluence in physioxia. Cells were trypsinized (0.25% Trypsin/EDTA; Corning), resuspended in basal chondrogenic media (93.24% High-Glucose DMEM (Gibco), 1% dexamethasone 10^−5^M (Sigma), 1% ITS+premix (Becton-Dickinson), 1% Glutamax (Hyclone), 1% 100 mM Sodium Pyruvate (Hyclone), 1% MEM Non-Essential Amino Acids (Hyclone), 0.26% L-Ascorbic Acid Phosphate 50mM (Wako), 0.5% Fungizone (Life Technologies) with TGF-β1 (Peprotech) and seeded as described below.

### Generation and Maintenance of 3D Aggregates

To generate 3D aggregates, cells were seeded at 50,000 cells per well (in 96-well cell repellent u-bottom plates, GreinerBio) and then centrifuged at 500 RCF, 5 min. For the TGF-β1 dose response studies, that serve as positive controls for the reporter cells, aggregates were cultured in basal chondrogenic media and different concentrations of TGF-β1 ranging from 0-10 ng/ml. For the DoE studies, aggregates were cultured in basal chondrogenic media (1ng/ml TGF-β1) as a control or basal media supplemented with vitamins and minerals (**Table 1** and **Supplemental Table 1**,**2**). Plates were cultured for three weeks in physioxia, media was sampled and replaced three times a week with respective medium. An OT-2 (Opentrons) python coded robotic pipette, programmed at an aspiration height of 2mm from the bottom of the wells and aspiration rate of 40μl/s was utilized for media preparation, cell feeding, and media sampling for luciferase assay **(Supplementary File 1)**. After three weeks, cell aggregates were either fixed in neutral buffered formalin for histology or medium removed and aggregates frozen dry (−20°C) for biochemical assays.

### Tissue Engineered Cartilage Sheets

#### A) Biochamber Sterilization and Assembly

Custom 3D printed biochambers^51^ that produce 1.2 cm^2^ cartilage sheets are shown in **Fig. 6**. The chambers are made of an acrylonitrile butadiene styrene (ABS) seeding chamber and a 10 μm pore polyester membrane (Sterlitech). Screws, a silicone washer and ABS frits hold everything securely and prevent any leaking in between the different pieces. Furthermore, they keep the chamber elevated to allow medium to reach the membrane from the top and bottom for efficient media exchange. The chambers are contained within Nalgene containers modified to have a 0.2 μm sterile filter on the top to allow gas exchange.

The Nalgene containers along with screws, silicone washer, polyester membrane and nuts were autoclaved and sterile filters fitted to the containers in a biosafety cabinet. ABS pieces were placed in a sealable container for sterilization by immersion in a 10% bleach solution, water rinse, followed by a 10% sodium thiosulfate treatment to neutralize any remaining chlorine, sterile water and isopropanol wash before drying in the biosafety cabinet. Biochambers were assembled as shown in **Fig. 6a** inside a biosafety hood using sterile surgical gloves and autoclaved surgical tools to handle biochamber parts. Once assembled, the polyester membrane was coated with fibronectin (50μg/cm^2^, Corning, in PBS) and allowed to dry in a biosafety cabinet for 1hr.

#### B) Generation and Culture of Tissue Engineered Cartilage Sheets

COL2A1-Gluc cells or uninfected primary rabbit chondrocytes were seeded at 5 × 10^6^ cells/cm^2^ in ABS biochambers with a 1.2 cm^2^ seeding area in basal chondrogenic media alone or in basal media with condition 25 at 1x and 1/15x (**Supplementary Table 2**)^51^. Media were added in the Nalgene container outside of the biochamber making sure it did not reach the top of the biochamber and combine with the cell suspension inside the biochamber in order to allow cells to adhere to the membrane. After 1 day, medium was added to the top of the biochamber so that media exchange occurs with the inside of the whole biochamber. These were cultured in physioxia on a shaker (10 RPM) for 3-weeks with media changes three times a week. During media replacement, samples from COL2A1-Gluc biochambers were assessed for luciferase. After three weeks, cartilage sheets were collected, four (4 mm) biopsy punches were taken for mechanical assessment, and collagen cross-linking analysis, remaining pieces of the sheets were frozen (−80°C) for biochemical assessment or stored in formalin for histology.

### Luciferase Assay

Cell culture medium sampled from the seeded 96-wells (20μL/well) was assessed using a stabilized *Gaussia* Luciferase buffer at a final concentration of 0.09 M MES, 0.15M Ascorbic Acid, and 4.2μM Coelenterazine in white 96-well plates. Luminescence was measured in a plate reader (25°C, relative light units, EnVision plate reader). An OT-2 (Opentrons) python coded robotic pipette was utilized for luciferase buffer addition to white plates (GreinerBio).

### Immunohistochemistry/ Immunofluorescence

At the end of three-week culture, cell aggregates were fixed in 10% Neutral Buffered Formalin overnight, embedded in paraffin wax and sectioned (7μm sections). Sections were deparaffinized and hydrated, followed by treatment with pronase (1mg/ml, Sigma P5147, in PBS with 5mM CaCl_2_) for epitope retrieval and incubation with primary anti-Collagen Type II (DSHB II-II6B3) followed by a biotinylated secondary and Streptavidin-HRP (BD Biosciences). II-II6B3 was deposited to the DSHB by Linsenmayer, T.F. (DSHB Hybridoma Product II-II6B3). Sections were stained with a chromogen-based substrate kit (Vector labs, VIP substrate vector kit). Engineered cartilage sheet sections were also treated with pronase and primary anti-Collagen Type II (DSHB II-II6B3) followed by VectaFluor R.T.U Antibody Kit DyLight® 488 (Vector Labs DI-2788) following manufacturer’s protocol. All sections were imaged at 10x magnification.

### Biochemical Assays

Frozen cell aggregates, or pieces of engineered cartilage were thawed in PBS, and enzymatically digested with Papain (25 μg/ml, Sigma, P4762, in 2mM cysteine; 50mM sodium phosphate; 2mM EDTA at a pH 6.5, 100 μl) at 65°C overnight. During digestion, plates were covered with a qPCR adhesive sealing film (USA Scientific), a silicone sheet, and steel plates clamped to the plate to prevent evaporation. After digestion half of the digest was transferred to another plate and frozen for hydroxyproline assessment. For the remaining half of the digest, papain was inactivated with 0.1M NaOH (50 μl) followed by neutralization (100mM Na2HPO4, 0.1 N HCL, pH 1.82, 50 μl). To assess DNA, samples of the digests (20 μl) were combined with buffered Hoechst dye (#33258, 667ng/ml, phosphate buffer pH 8, 100 μl) and fluorescence measured at an excitation of 365nm and emission of 460nm. For GAG assessment, samples of aggregate digest (5μl) were combined with a 1,9-Dimethyl-methylene blue solution (195μl) and absorbance was measured at 595nm and 525nm^62^. Absorbance readings were corrected by subtracting 595nm reading from 525nm. Total micrograms of DNA and GAG were calculated using a Calf thymus DNA standard (Sigma) and Chondroitin Sulfate standard (Seikagaku Corp.), respectively.

For hydroxyproline (HP), the frozen digest was thawed and incubated overnight at 105°C with 6M HCL (200μl). Plates were covered as described above to prevent evaporation. Samples were subsequently dried at 70°C overnight with a hydroxyproline standard (Sigma). Copper sulfate (0.15M, 10μl) and sodium hydroxide (2.5M, 10μl) were added to each well and incubated at 50°C for 5 minutes, followed by hydrogen peroxide (6%, 10 μl) for 10minutes. Sulfuric acid (1.5 M, 40μl) and Ehrlich’s reagent^21^ (20μl) were added and samples further incubated at 70°C for 15 minutes before reading absorbance at 505nm. Total micrograms of hydroxyproline were calculated using the standard. Total collagen was calculated by the following formula^21^ (μg of HP X 7.6 = μg Total Collagen).^21^

### Collagen Cross-Link Analysis

After harvest, samples were frozen at -80°C until use. Samples were lyophilized, and dry weights obtained. Proteoglycans were extracted using 4M guanidine hydrochloride (GuHCl) in 50mM Tris buffer pH 7.4. The residue was exhaustively rinsed using MilliQ water to remove residual GuHCl, lyophilized and weighed. The cross-linked collagen network was depolymerized using equal volumes of pepsin (0.5mg/mL in 0.5M acetic acid)^26^. Equivalent aliquots of dry weights were loaded on 6% polyacrylamide gels. Pepsin-extracted type II collagen from adult rabbit articular cartilage was used as a control. After electrophoresis, collagen chains were stained using Coomassie Blue. For Western blots, following SDS-PAGE the separated collagen chains were transferred, by electrophoresis, onto a polyvinyl difluoride (PVDF) membrane and probed with monoclonal antibody (mAb) 10F2 to identify the C-telopeptide of α1(II) collagen chains cross-linked to α1(II) chains. Another blot was probed with polyclonal antibody (pAb) 5890 to identify the N-telopeptide of α1(XI) chains cross-linked to α1(II) chains. This blot was then stripped and probed with mAb 1C10 to identify α1(II) chains. As we have described before, this determines if a heteropolymer of type II and type XI collagen had formed^26^.

### Mechanical Testing

#### A) Compression Testing

Biopsy punches (4mm) were thawed in Tyrode’s solution (Sigma) with protease inhibitors (Sigma, P8465) and equilibrated to room temperature. Using a TA.XT*Plus connect* Texture analyzer a trigger force of 0.1 gram determined the height of the tissue, then 5-20% strain was applied in 5% increments with a 20-minute hold to reach equilibrium. From these results, the equilibrium force was calculated, and a stress vs strain curve generated^21^. Young’s modulus in compression was determined from the slope of these curves.

#### B) Tensile Testing

Biopsy punches (4mm) were thawed in Tyrode’s solution (Sigma) with protease inhibitors and equilibrated to room temperature. As previously described, a custom dog-bone punch was made from biopsy punches and punches taken from the 4 mm punch^21^. Custom holders were made from laminate projector sheets and dog-bone punches attached using cyanoacrylate glue (Ultra Gel Control, Loctite), tissue was continuously bathed in PBS during this process^21^. Using a TA.XTPlus connect Texture analyzer with a trigger force of 0.1 gram, tissues were stretched to failure, the equilibrium tensile force was calculated and a stress vs strain curve generated^21^. Young’s modulus in tension was determined from the slope of these curves.

### Design of Experiment Response Surface Design

Design-Expert 12 (StatEase) was used to generate a surface response model to assess the effect of 15 factors: linoleic acid, cobalt, copper, chromium, iodine, manganese, molybdenum, thyroxine, vitamin A, vitamin B12, vitamin B7, vitamin D, vitamin E, vitamin K, and zinc. **Table 1** shows minimum and maximum concentrations input into Design-Expert. The response surface I-optimal blocked design generated 240 total conditions to assess the response (**Supplemental Table 1**).

### Design of Experiments Analysis

At the end of this experiment, responses (luminescence, metabolic activity and aggregate area) from the screen of 240 conditions as well as results from the one factor at a time approach, were analyzed (Design-Expert, StatEase). Analysis of the results suggested a quadratic model as the best fit. After transformation of the data to fit a quadratic model, Analysis of Variance (ANOVA) was used to identify the positive and negative effects on chondrogenesis as well as fit statistics for the model. The optimization module of the software was used to generate five predicted optimal combinations of factors (**Supplemental Table 2**). Two sets of parameters were used to generate predicted conditions. For condition 25 from **Supplemental Table 2**, all vitamins and minerals were targeted at 75% serum max except for linolenic acid, vitamin A, copper and vitamin D which are set at their predicted optima. For the other predicted conditions vitamins and minerals were set between 0.01% of serum max and serum max except for vitamin A, E and linolenic acid which were at their approximate optima. All conditions were selected to maximize luminescence for week two and week three as well as aggregate area for week three. Condition 25 also had a target of 0.2, for Resazurin (metabolic activity), the average measurement for chondrocyte aggregates, at week three.

### Statistical Analysis

Statistical analysis for all experiments except for the Design of Experiments screen (analysis described above) were performed using GraphPad Prism 9 and One-way or two-way ANOVA. All data passed tests for normality. In all figures * indicates p-value < 0.05, ** indicates p-value <0.01, and *** indicates p-value < 0.001.

## Supporting information

other supplemental

## ACKNOWLEDGEMENTS

We would like to thank Dr. Steven Mills (University of Texas Health Sciences) for the donation of rabbit tissue. The plasmids, pMD2.G, and psPAX2, were a gift from Didier Trono (Addgene plasmid # 12259; http://n2t.net/addgene:12259; RRID:Addgene_12259) and (Addgene plasmid # 12260; http://n2t.net/addgene:12260; RRID:Addgene_12260) respectively. Funding was supplied by the University of Central Florida (TJK), University of Central Florida College of Medicine (TJK), the Rolanette and Berdon Lawrence Bone Disease Program (TJK) and NIH grant AR057025 (RFJ).

