## Supplemental Table 1 for "Micronutrient Optimization Using Design of Experiments Approach in Tissue Engineered Articular Cartilage for Production of Type II Collagen"

| Runs (Conditions) | ALA | Cr | Co | Cu | I | Mn | Concent<br>Mo |
|---|---|---|---|---|---|---|---|
| 1 | 13.5 | 0.028 | 0.000171 | 0 | 0 | 0 | 0 |
| 2 | 7.425 | 0 | 0.00045 | 0.67 | 0 | 0 | 0.001 |
| 3 | 13.5 | 0.028 | 0.00045 | 0.67 | 0.046 | 0 | 0 |
| 4 | 13.5 | 0 | 0 | 0.67 | 0 | 0.036 | 0 |
| 5 | 13.5 | 0 | 0 | 0 | 0 | 0.036 | 0.001 |
| 6 | 13.5 | 0 | 0.00045 | 0 | 0.046 | 0.0207 | 0 |
| 7 | 13.5 | 0 | 0.00045 | 0 | 0 | 0 | 0.001 |
| 8 | 0 | 0.028 | 0.00045 | 0 | 0.046 | 0 | 0.001 |
| 9 | 13.5 | 0.01624 | 0.000304 | 0.67 | 0.034725 | 0.036 | 0.00091 |
| 10 | 0 | 0 | 0.00045 | 0 | 0.046 | 0.036 | 0 |
| 11 | 0 | 0 | 0 | 0.67 | 0 | 0 | 0.001 |
| 12 | 13.5 | 0 | 0 | 0.3484 | 0.046 | 0.019694 | 0 |
| 13 | 13.5 | 0.028 | 0 | 0 | 0 | 0.036 | 0.001 |
| 14 | 0 | 0 | 0.00045 | 0 | 0.046 | 0 | 0.001 |
| 15 | 13.5 | 0.028 | 0.00045 | 0 | 0 | 0.036 | 0 |
| 16 | 13.5 | 0 | 0.00045 | 0 | 0 | 0.036 | 0 |
| 17 | 13.5 | 0.028 | 0.00045 | 0.67 | 0 | 0 | 0 |
| 18 | 13.5 | 0.028 | 0.000158 | 0.67 | 0.046 | 0 | 0.000515 |
| 19 | 5.0625 | 0.028 | 0 | 0 | 0.046 | 0 | 0.001 |
| 20 | 13.5 | 0.028 | 0 | 0.67 | 0.046 | 0 | 0 |
| 21 | 0 | 0.01218 | 0.00045 | 0.67 | 0 | 0.036 | 0.001 |
| 22 | 0 | 0.028 | 0.00045 | 0 | 0.046 | 0.036 | 0 |
| 23 | 0 | 0.028 | 0.00045 | 0.67 | 0.046 | 0.036 | 0 |
| 24 | 0 | 0.028 | 0.00045 | 0 | 0 | 0.036 | 0.00041 |
| 25 | 6.2775 | 0.028 | 0 | 0 | 0 | 0.036 | 0.001 |
| 26 | 13.5 | 0 | 0 | 0 | 0.046 | 0.036 | 0.001 |
| 27 | 13.5 | 0 | 0 | 0.67 | 0 | 0 | 0 |
| 28 | 9.23046 | 0.028 | 3.38E-05 | 0.67 | 0.04485 | 0.008491 | 0.000699 |
| 29 | 7.155 | 0 | 0.000236 | 0.67 | 0.046 | 0.036 | 0.001 |
| 30 | 13.5 | 0 | 0 | 0 | 0 | 0 | 0 |
| 31 | 0 | 0 | 0 | 0 | 0 | 0 | 0 |
| 32 | 0 | 0.028 | 0 | 0.67 | 0.046 | 0 | 0.001 |
| 33 | 0 | 0 | 0.00045 | 0 | 0.027813 | 0.036 | 0.001 |
| 34 | 0 | 0.028 | 0 | 0.67 | 0.0184 | 0.036 | 0.001 |
| 35 | 8.91829 | 0.00112 | 9.06E-05 | 0.67 | 0.024334 | 0.01386 | 0.000705 |
| 36 | 0 | 0.01456 | 0.000338 | 0.67 | 0.02346 | 0.00936 | 0.00007 |
| 37 | 13.5 | 0.028 | 0.00045 | 0.3685 | 0.046 | 0.036 | 0.001 |
| 38 | 0 | 0.0147 | 0 | 0.67 | 0 | 0.036 | 0 |
| 39 | 13.5 | 0 | 0.00045 | 0 | 0.0253 | 0 | 0.00056 |
| 40 | 13.5 | 0.028 | 0.00045 | 0 | 0 | 0 | 0.001 |
| 41 | 0 | 0 | 0 | 0 | 0 | 0 | 0 |
| 42 | 5.67 | 0.028 | 0 | 0.67 | 0.046 | 0.036 | 0 |
| 43 | 0 | 0.028 | 0 | 0 | 0.046 | 0.036 | 0 |
| 44 | 0 | 0.028 | 0 | 0.67 | 0 | 0 | 0.001 |

| | | | | | | | |
|---|---|---|---|---|---|---|---|
| 45 | 0 | 0 | 0 | 0.67 | 0 | 0.036 | 0.001 |
| 46 | 0 | 0 | 0.00045 | 0 | 0 | 0 | 0.001 |
| 47 | 13.5 | 0 | 0.000254 | 0.35175 | 0.046 | 0.036 | 0.001 |
| 48 | 0 | 0 | 0 | 0 | 0 | 0 | 0 |
| 49 | 0 | 0 | 0.00045 | 0.67 | 0.046 | 0 | 0.000425 |
| 50 | 13.5 | 0.0147 | 0.00045 | 0.67 | 0.046 | 0.036 | 0.001 |
| 51 | 0 | 0 | 0.00045 | 0.67 | 0 | 0.036 | 0 |
| 52 | 0 | 0 | 0.00045 | 0.36515 | 0.046 | 0 | 0 |
| 53 | 0 | 0.028 | 0.00045 | 0.67 | 0 | 0.036 | 0.001 |
| 54 | 0 | 0 | 0.00045 | 0.67 | 0.046 | 0.036 | 0.001 |
| 55 | 13.5 | 0.0161 | 0 | 0.67 | 0 | 0 | 0.001 |
| 56 | 0 | 0 | 0 | 0 | 0 | 0 | 0 |
| 57 | 0 | 0 | 0 | 0 | 0.046 | 0.036 | 0 |
| 58 | 13.5 | 0.028 | 0 | 0 | 0.046 | 0 | 0.001 |
| 59 | 0 | 0.028 | 0 | 0 | 0.046 | 0 | 0 |
| 60 | 10.125 | 0.028 | 0.00045 | 0.67 | 0 | 0.0216 | 0 |
| 61 | 7.695 | 0.028 | 0.00045 | 0 | 0 | 0.036 | 0.001 |
| 62 | 0 | 0 | 0 | 0.67 | 0 | 0.036 | 0 |
| 63 | 0 | 0 | 0 | 0 | 0 | 0 | 0 |
| 64 | 0 | 0 | 0.000191 | 0.67 | 0 | 0.0198 | 0.001 |
| 65 | 13.5 | 0.028 | 0 | 0.67 | 0 | 0.01152 | 0.001 |
| 66 | 13.5 | 0 | 0 | 0 | 0.046 | 0 | 0.001 |
| 67 | 0 | 0 | 0.000225 | 0.67 | 0.046 | 0 | 0.001 |
| 68 | 13.5 | 0 | 0 | 0.67 | 0.046 | 0.036 | 0 |
| 69 | 8.775 | 0.028 | 0 | 0 | 0 | 0.0234 | 0 |
| 70 | 0 | 0 | 0.00045 | 0.3417 | 0 | 0 | 0 |
| 71 | 0 | 0 | 0.00045 | 0 | 0 | 0.036 | 0 |
| 72 | 0 | 0.028 | 0.00045 | 0.67 | 0.046 | 0.036 | 0 |
| 73 | 13.5 | 0 | 0.00045 | 0.67 | 0.046 | 0.036 | 0 |
| 74 | 13.5 | 0.028 | 0.00045 | 0 | 0 | 0 | 0.001 |
| 75 | 6.4125 | 0.028 | 0.00045 | 0.67 | 0.046 | 0 | 0.001 |
| 76 | 13.5 | 0.028 | 0.00045 | 0.67 | 0 | 0.036 | 0.001 |
| 77 | 0 | 0.028 | 0 | 0 | 0 | 0.01656 | 0 |
| 78 | 0 | 0.028 | 0.000448 | 0.49915 | 0.034165 | 0.01458 | 0.000821 |
| 79 | 0 | 0 | 0.00045 | 0 | 0 | 0 | 0.00048 |
| 80 | 13.5 | 0 | 0.00045 | 0.67 | 0 | 0 | 0.001 |
| 81 | 13.5 | 0.028 | 0 | 0 | 0 | 0.036 | 0 |
| 82 | 0 | 0.028 | 0.00045 | 0 | 0.046 | 0 | 0 |
| 83 | 13.5 | 0 | 0 | 0 | 0.046 | 0.036 | 0.000485 |
| 84 | 0 | 0.028 | 0 | 0 | 0 | 0.036 | 0.00052 |
| 85 | 13.5 | 0 | 0 | 0.67 | 0.01817 | 0.036 | 0.001 |
| 86 | 13.5 | 0 | 0 | 0.67 | 0.046 | 0.036 | 0.001 |
| 87 | 13.5 | 0.028 | 0.00045 | 0.67 | 0.01932 | 0.036 | 0 |
| 88 | 0 | 0.028 | 0 | 0.477405 | 0.045892 | 0 | 0.000338 |
| 89 | 13.5 | 0 | 0 | 0.67 | 0 | 0.02124 | 0.001 |
| 90 | 7.81988 | 0.01022 | 0.00018 | 0.0335 | 0.04071 | 0 | 0.000671 |
| 91 | 13.5 | 0 | 0.00045 | 0.67 | 0 | 0 | 0 |

| | | | | | | | |
|---|---|---|---|---|---|---|---|
| 92 | 0 | 0 | 0 | 0.67 | 0 | 0 | 0 |
| 93 | 0 | 0.028 | 0.000189 | 0.67 | 0.046 | 0.036 | 0 |
| 94 | 13.5 | 0 | 0.00045 | 0.35175 | 0 | 0.036 | 0.001 |
| 95 | 0 | 0.01414 | 0 | 0 | 0.046 | 0.036 | 0.001 |
| 96 | 0 | 0.028 | 0 | 0 | 0.01725 | 0.036 | 0.001 |
| 97 | 13.5 | 0.028 | 0.00045 | 0 | 0 | 0.036 | 0 |
| 98 | 0 | 0.0112 | 0 | 0 | 0 | 0 | 0.001 |
| 99 | 0 | 0 | 0.00045 | 0 | 0 | 0.0144 | 0 |
| 100 | 13.5 | 0.0126 | 0.00045 | 0.67 | 0.046 | 0 | 0.001 |
| 101 | 13.5 | 0 | 0 | 0.67 | 0.046 | 0 | 0 |
| 102 | 13.5 | 0.028 | 0 | 0.67 | 0.046 | 0 | 0 |
| 103 | 13.5 | 0.028 | 0.00045 | 0 | 0 | 0.036 | 0.001 |
| 104 | 13.5 | 0 | 0.000185 | 0 | 0.046 | 0 | 0.001 |
| 105 | 6.5475 | 0 | 0.00045 | 0 | 0.046 | 0.036 | 0.000565 |
| 106 | 13.5 | 0.028 | 0 | 0.67 | 0 | 0 | 0 |
| 107 | 0 | 0 | 0 | 0 | 0.046 | 0 | 0.00045 |
| 108 | 13.5 | 0.028 | 0 | 0 | 0 | 0.036 | 0.001 |
| 109 | 13.5 | 0.028 | 0.00045 | 0 | 0.046 | 0 | 0.001 |
| 110 | 0 | 0.028 | 0.00045 | 0.67 | 0.046 | 0.036 | 0.001 |
| 111 | 13.5 | 0.028 | 0 | 0 | 0 | 0 | 0.001 |
| 112 | 0 | 0.0154 | 0 | 0.67 | 0 | 0.036 | 0.001 |
| 113 | 0 | 0 | 0.00045 | 0.67 | 0 | 0 | 0.001 |
| 114 | 0 | 0 | 0 | 0 | 0 | 0 | 0 |
| 115 | 0 | 0 | 0 | 0 | 0 | 0 | 0 |
| 116 | 0 | 0.028 | 0.00045 | 0.67 | 0 | 0 | 0 |
| 117 | 5.13 | 0.028 | 0 | 0.33835 | 0.046 | 0 | 0 |
| 118 | 0 | 0 | 0 | 0 | 0.046 | 0.036 | 0.001 |
| 119 | 13.5 | 0 | 0.00045 | 0 | 0.046 | 0 | 0 |
| 120 | 13.5 | 0.0147 | 0.00045 | 0 | 0 | 0.036 | 0 |
| 121 | 13.5 | 0 | 0 | 0 | 0 | 0 | 0.00041 |
| 122 | 7.155 | 0.0154 | 0.00045 | 0 | 0 | 0 | 0.0005 |
| 123 | 6.25718 | 0.028 | 0.00045 | 0.3015 | 0.046 | 0.036 | 0 |
| 124 | 13.5 | 0 | 0.000263 | 0.67 | 0 | 0.036 | 0 |
| 125 | 13.5 | 0.028 | 0 | 0.67 | 0.046 | 0.036 | 0 |
| 126 | 13.5 | 0.028 | 0 | 0.67 | 0 | 0 | 0 |
| 127 | 0 | 0 | 0 | 0.67 | 0.046 | 0 | 0.001 |
| 128 | 13.5 | 0 | 0 | 0.67 | 0.046 | 0 | 0.001 |
| 129 | 13.5 | 0 | 0.00045 | 0 | 0 | 0 | 0 |
| 130 | 13.5 | 0.01484 | 0 | 0 | 0 | 0 | 0.001 |
| 131 | 0 | 0.028 | 0.00045 | 0.32495 | 0.02507 | 0 | 0.001 |
| 132 | 13.5 | 0 | 0.00045 | 0 | 0.046 | 0 | 0.001 |
| 133 | 0 | 0.028 | 0 | 0 | 0 | 0 | 0 |
| 134 | 13.5 | 0.028 | 0.00045 | 0.27135 | 0 | 0 | 0 |
| 135 | 0 | 0 | 0 | 0.67 | 0 | 0 | 0.001 |
| 136 | 0 | 0 | 0 | 0 | 0 | 0 | 0 |
| 137 | 13.5 | 0.0126 | 0.00045 | 0.31155 | 0.0253 | 0.018 | 0.000425 |
| 138 | 0 | 0.028 | 0 | 0.67 | 0 | 0 | 0 |

| | | | | | | | |
|---|---|---|---|---|---|---|---|
| 139 | 13.5 | 0.028 | 0 | 0.67 | 0.046 | 0.036 | 0.001 |
| 140 | 13.5 | 0.028 | 0.00045 | 0 | 0.046 | 0.036 | 0.001 |
| 141 | 0 | 0.028 | 0 | 0 | 0.046 | 0.01944 | 0.001 |
| 142 | 0 | 0.028 | 0.00045 | 0 | 0 | 0 | 0 |
| 143 | 0 | 0.014687 | 0.000268 | 0 | 0.023 | 0.036 | 0 |
| 144 | 0 | 0.028 | 0.00045 | 0 | 0 | 0.036 | 0.001 |
| 145 | 7.425 | 0 | 0 | 0.67 | 0.02645 | 0.036 | 0 |
| 146 | 13.5 | 0.028 | 0.000239 | 0.67 | 0 | 0.036 | 0.001 |
| 147 | 13.5 | 0 | 0.00045 | 0 | 0.046 | 0 | 0 |
| 148 | 0 | 0.028 | 0 | 0 | 0.046 | 0.036 | 0.001 |
| 149 | 0 | 0.028 | 0 | 0.67 | 0.046 | 0.036 | 0.001 |
| 150 | 0 | 0.028 | 0.000203 | 0.67 | 0 | 0 | 0.001 |
| 151 | 13.5 | 0.028 | 0 | 0 | 0.046 | 0.036 | 0 |
| 152 | 0 | 0 | 0 | 0.67 | 0.01955 | 0 | 0 |
| 153 | 0 | 0.028 | 0.00045 | 0.67 | 0.046 | 0 | 0 |
| 154 | 13.5 | 0.028 | 0.00045 | 0 | 0.046 | 0 | 0 |
| 155 | 13.5 | 0.028 | 0.000225 | 0.67 | 0 | 0.036 | 0.001 |
| 156 | 0 | 0.028 | 0.00045 | 0.67 | 0.046 | 0 | 0.00058 |
| 157 | 13.5 | 0 | 0 | 0 | 0 | 0.036 | 0.001 |
| 158 | 13.5 | 0.028 | 0 | 0.67 | 0.046 | 0 | 0.001 |
| 159 | 13.5 | 0.028 | 0.00045 | 0 | 0 | 0 | 0 |
| 160 | 0 | 0.028 | 0.00045 | 0 | 0.046 | 0.036 | 0.001 |
| 161 | 0 | 0 | 0.00045 | 0.67 | 0.046 | 0.036 | 0 |
| 162 | 0 | 0.028 | 0 | 0.416993 | 0.046 | 0 | 0 |
| 163 | 0 | 0 | 0 | 0 | 0 | 0 | 0 |
| 164 | 13.5 | 0 | 0 | 0.3149 | 0.046 | 0 | 0.001 |
| 165 | 0 | 0 | 0.00045 | 0.67 | 0 | 0.036 | 0 |
| 166 | 0 | 0 | 0 | 0 | 0 | 0 | 0 |
| 167 | 13.5 | 0 | 0.00045 | 0.67 | 0.046 | 0 | 0.001 |
| 168 | 13.5 | 0.028 | 0.00045 | 0.67 | 0.046 | 0 | 0.001 |
| 169 | 0 | 0 | 0.00045 | 0 | 0.046 | 0.018 | 0.001 |
| 170 | 0 | 0 | 0 | 0 | 0 | 0.01854 | 0 |
| 171 | 1.9575 | 0.02618 | 0.00045 | 0.22457 | 0 | 0.03132 | 0.000455 |
| 172 | 13.5 | 0.028 | 0 | 0.67 | 0 | 0.036 | 0 |
| 173 | 0 | 0.028 | 0.00045 | 0.67 | 0 | 0.036 | 0.001 |
| 174 | 0 | 0 | 0.000248 | 0 | 0 | 0 | 0.001 |
| 175 | 13.5 | 0 | 0 | 0 | 0.02392 | 0.036 | 0 |
| 176 | 0 | 0 | 0 | 0.67 | 0.046 | 0.036 | 0 |
| 177 | 0 | 0 | 0 | 0 | 0.046 | 0 | 0 |
| 178 | 0 | 0 | 0 | 0 | 0 | 0.036 | 0.001 |
| 179 | 0 | 0.028 | 0.00045 | 0 | 0 | 0 | 0.001 |
| 180 | 13.5 | 0 | 0.00045 | 0.67 | 0 | 0.036 | 0.00046 |
| 181 | 0 | 0.028 | 0.000218 | 0 | 0 | 0 | 0.001 |
| 182 | 13.5 | 0.028 | 0 | 0.67 | 0 | 0 | 0.001 |
| 183 | 6.4125 | 0 | 0.00045 | 0 | 0.046 | 0 | 0.00064 |
| 184 | 13.5 | 0 | 0.00045 | 0.67 | 0.046 | 0 | 0.001 |
| 185 | 13.5 | 0 | 0 | 0 | 0 | 0 | 0 |

| | | | | | | | |
|---|---|---|---|---|---|---|---|
| 186 | 0 | 0 | 0 | 0.67 | 0.01932 | 0 | 0 |
| 187 | 0 | 0.028 | 0 | 0.67 | 0.046 | 0.036 | 0.001 |
| 188 | 0 | 0 | 0 | 0 | 0 | 0 | 0 |
| 189 | 13.5 | 0.028 | 0.00045 | 0.67 | 0.046 | 0 | 0 |
| 190 | 13.5 | 0.028 | 0.00045 | 0 | 0.046 | 0.036 | 0 |
| 191 | 0 | 0 | 0 | 0 | 0 | 0.036 | 0.001 |
| 192 | 0 | 0 | 0 | 0 | 0 | 0 | 0 |
| 193 | 0 | 0 | 0 | 0 | 0.046 | 0.036 | 0 |
| 194 | 3.8804 | 0.016788 | 0.00041 | 0 | 0.00391 | 0.026038 | 0 |
| 195 | 6.21 | 0.028 | 0 | 0.67 | 0 | 0 | 0.0005 |
| 196 | 0 | 0.028 | 0.000203 | 0.67 | 0 | 0 | 0 |
| 197 | 6.75 | 0.028 | 0 | 0 | 0.02415 | 0 | 0 |
| 198 | 13.5 | 0 | 0.00045 | 0 | 0 | 0.036 | 0.001 |
| 199 | 4.05055 | 0.02674 | 7.33E-05 | 0.56615 | 0.03473 | 0.01302 | 0.001 |
| 200 | 0 | 0 | 0 | 0 | 0 | 0 | 0 |
| 201 | 13.5 | 0 | 0.00045 | 0.67 | 0.046 | 0.036 | 0 |
| 202 | 0 | 0 | 0.00045 | 0.67 | 0 | 0 | 0.001 |
| 203 | 13.5 | 0 | 0 | 0 | 0 | 0 | 0 |
| 204 | 13.5 | 0.0154 | 0 | 0 | 0.046 | 0.036 | 0 |
| 205 | 8.5725 | 0.028 | 0.000245 | 0.67 | 0.046 | 0.01782 | 0.001 |
| 206 | 0 | 0 | 0.00045 | 0 | 0.046 | 0 | 0.001 |
| 207 | 0 | 0 | 0 | 0.67 | 0.046 | 0.036 | 0.001 |
| 208 | 13.5 | 0.028 | 0 | 0 | 0.046 | 0 | 0.0004 |
| 209 | 0 | 0 | 0.00045 | 0.67 | 0 | 0.036 | 0.001 |
| 210 | 13.5 | 0 | 0 | 0 | 0.046 | 0.036 | 0.001 |
| 211 | 0 | 0 | 0 | 0 | 0 | 0 | 0.001 |
| 212 | 13.5 | 0 | 0 | 0.67 | 0 | 0 | 0 |
| 213 | 13.5 | 0.028 | 0.00045 | 0.67 | 0 | 0.036 | 0 |
| 214 | 13.5 | 0 | 0 | 0.67 | 0.046 | 0.036 | 0.001 |
| 215 | 13.5 | 0 | 0.00045 | 0 | 0 | 0.036 | 0.001 |
| 216 | 13.2975 | 0.0098 | 0 | 0.414682 | 0.00023 | 0.014963 | 0.000662 |
| 217 | 0 | 0.028 | 0.00045 | 0 | 0.046 | 0 | 0.001 |
| 218 | 0 | 0 | 0.00045 | 0.67 | 0 | 0.036 | 0 |
| 219 | 13.5 | 0.028 | 0 | 0 | 0.046 | 0.036 | 0.001 |
| 220 | 13.5 | 0.028 | 0 | 0.3685 | 0 | 0.036 | 0 |
| 221 | 13.5 | 0.01246 | 0.00045 | 0 | 0.046 | 0.01638 | 0.001 |
| 222 | 0 | 0.028 | 0.00045 | 0.335 | 0 | 0.036 | 0 |
| 223 | 0 | 0 | 0.00045 | 0 | 0.01955 | 0.036 | 0 |
| 224 | 0 | 0.028 | 0.00045 | 0.67 | 0.046 | 0.036 | 0.001 |
| 225 | 5.01935 | 0.028 | 0 | 0 | 0.046 | 0 | 0 |
| 226 | 13.5 | 0 | 0.00045 | 0.3484 | 0.046 | 0.036 | 0 |
| 227 | 0 | 0.028 | 0 | 0 | 0.02047 | 0.036 | 0 |
| 228 | 0 | 0.01064 | 0.00045 | 0.67 | 0.046 | 0 | 0 |
| 229 | 13.5 | 0.028 | 0.00045 | 0 | 0 | 0.018 | 0.001 |
| 230 | 0 | 0.028 | 0.00045 | 0.67 | 0 | 0 | 0 |
| 231 | 0 | 0.028 | 0.00045 | 0 | 0.046 | 0 | 0 |
| 232 | 13.5 | 0 | 0.00045 | 0.67 | 0.046 | 0.036 | 0.0004 |

| | | | | | | | |
|---|---|---|---|---|---|---|---|
| 233 | 0 | 0 | 0 | 0 | 0.046 | 0.036 | 0 |
| 234 | 13.5 | 0.028 | 0.00045 | 0 | 0.046 | 0 | 0.001 |
| 235 | 13.5 | 0 | 0.00045 | 0.67 | 0 | 0 | 0 |
| 236 | 13.5 | 0.028 | 0.00045 | 0.67 | 0 | 0.036 | 0.001 |
| 237 | 0 | 0.028 | 0 | 0.67 | 0 | 0.036 | 0 |
| 238 | 13.5 | 0.02044 | 0.000218 | 0.45895 | 0.01771 | 0 | 0 |
| 239 | 0 | 0 | 0 | 0.67 | 0.046 | 0 | 0 |
| 240 | 0 | 0.028 | 0.00045 | 0 | 0 | 0.036 | 0.00035 |

**ration (ug/ml)**

| T4 | Vit A | Vit B12 | Vit B7 | Vit D | Vit E | Vit K | Zn |
|---|---|---|---|---|---|---|---|
| 0 | 0 | 0.000914 | 0.005257 | 0 | 18.4 | 0 | 2.7 |
| 0 | 0 | 0.000914 | 0.005257 | 0.000301 | 0 | 0 | 3 |
| 0 | 1.56E-06 | 0 | 0 | 0.000301 | 18.4 | 0.0011 | 6 |
| 0.125 | 1.56E-06 | 0 | 0.005257 | 0.000301 | 0 | 0.0011 | 0 |
| 0 | 0 | 0.000914 | 0.005257 | 0.000301 | 18.4 | 0.0011 | 6 |
| 0 | 1.56E-06 | 0.000914 | 0.002734 | 0 | 18.4 | 0 | 6 |
| 0.125 | 1.56E-06 | 0.000914 | 0 | 0.000301 | 0 | 0 | 0 |
| 0.125 | 1.56E-06 | 0 | 0 | 0 | 18.4 | 0.0011 | 0 |
| 0.095 | 0 | 0 | 0.001551 | 7.53E-05 | 13.156 | 0.000578 | 6 |
| 0 | 1.56E-06 | 0.000914 | 0.005257 | 0 | 0 | 0 | 0 |
| 0 | 1.56E-06 | 0 | 0.005257 | 0.000301 | 18.4 | 0.0011 | 0 |
| 0.125 | 1.56E-06 | 0 | 0 | 0.000135 | 0 | 0 | 6 |
| 0.125 | 0 | 0.000914 | 0.005257 | 0 | 18.4 | 0.0011 | 0 |
| 0 | 1.56E-06 | 0.000914 | 0 | 0.000301 | 18.4 | 0.0011 | 6 |
| 0 | 0 | 0 | 0 | 0.000301 | 18.4 | 0 | 6 |
| 0.125 | 0 | 0.000914 | 0 | 0 | 18.4 | 0 | 0 |
| 0.125 | 1.56E-06 | 0.000914 | 0 | 0 | 18.4 | 0.0011 | 0 |
| 0 | 6.63E-07 | 0.000914 | 0.005257 | 0.000301 | 0 | 0 | 6 |
| 0 | 1.56E-06 | 0.000471 | 0.005257 | 0 | 18.4 | 0 | 0 |
| 0.125 | 1.56E-06 | 0.000914 | 0.005257 | 0.000301 | 9.2 | 0.0011 | 0 |
| 0 | 0 | 0.000377 | 0.00368 | 0 | 0 | 0.0011 | 0 |
| 0 | 0 | 0 | 0 | 0.000173 | 0 | 0.0011 | 0 |
| 0.125 | 0 | 0 | 0.005257 | 0.000301 | 0 | 0 | 6 |
| 0 | 1.56E-06 | 0.000914 | 0 | 0.000301 | 0 | 0 | 6 |
| 0.125 | 1.56E-06 | 0.000452 | 0.005257 | 0.000188 | 0 | 0.0011 | 6 |
| 0.0675 | 0 | 0 | 0.005257 | 0.000301 | 0 | 0.0011 | 0 |
| 0.125 | 0 | 0.000914 | 0 | 0.000301 | 18.4 | 0 | 0 |
| 0.125 | 0 | 0.000914 | 0.003295 | 0 | 0 | 0.00099 | 6 |
| 0 | 0 | 0.000914 | 0 | 0.000301 | 18.4 | 0 | 6 |
| 0 | 1.56E-06 | 0 | 0 | 0.000301 | 0 | 0.0011 | 3.6 |
| 0.125 | 0 | 0 | 0 | 0 | 0 | 0.000523 | 6 |
| 0 | 0 | 0.000914 | 0.002497 | 0 | 8.832 | 0.000385 | 0 |
| 0.125 | 1.56E-06 | 0.000914 | 0.005257 | 0.000134 | 18.4 | 0.0011 | 0 |
| 0 | 6.63E-07 | 0.000914 | 0 | 0.000301 | 8.004 | 0.0011 | 0 |
| 0.0025 | 1.48E-06 | 0.000428 | 0.003944 | 6.96E-05 | 9.79041 | 0.000127 | 5.8768 |
| 0.125 | 0 | 0.000361 | 0.004705 | 0.000224 | 18.4 | 0.0011 | 6 |
| 0.053125 | 9.94E-07 | 0 | 0 | 0 | 0 | 0 | 0 |
| 0 | 1.56E-06 | 0.000914 | 0.005257 | 0 | 18.4 | 0 | 0 |
| 0 | 0 | 0 | 0 | 0.000301 | 18.4 | 0.0011 | 0 |
| 0 | 0 | 0.000914 | 0 | 0 | 0 | 0.0011 | 6 |
| 0 | 0 | 0 | 0 | 0 | 0 | 0 | 0 |
| 0 | 1.56E-06 | 0.000567 | 0 | 0 | 8.372 | 0 | 6 |
| 0.125 | 0 | 0.000914 | 0 | 0.000301 | 18.4 | 0 | 6 |
| 0.061144 | 1.56E-06 | 0 | 0 | 0 | 0 | 0.0011 | 6 |

| | | | | | | | |
|---|---|---|---|---|---|---|---|
| 0 | 0 | 0 | 0 | 0 | 18.4 | 0 | 3.15 |
| 0.125 | 1.56E-06 | 0 | 0.005257 | 0.000301 | 0 | 0 | 6 |
| 0 | 1.56E-06 | 0 | 0.005257 | 0.000301 | 18.4 | 0 | 0 |
| 0 | 0 | 0 | 0 | 0 | 0 | 0 | 0 |
| 0.125 | 0 | 0.000914 | 0 | 0.000301 | 0 | 0.0011 | 0 |
| 0.125 | 1.56E-06 | 0.000914 | 0 | 0 | 18.4 | 0.0011 | 6 |
| 0.125 | 1.56E-06 | 0 | 0.005257 | 0 | 18.4 | 0.0011 | 6 |
| 0 | 0 | 0 | 0.005257 | 0 | 9.936 | 0.0011 | 3.45 |
| 0.125 | 1.56E-06 | 0 | 0 | 0.000301 | 18.4 | 0 | 0 |
| 0.125 | 0 | 0.000914 | 0.002155 | 0 | 0 | 0 | 6 |
| 0.125 | 0 | 0 | 0.005257 | 0 | 0 | 0 | 6 |
| 0 | 0 | 0 | 0 | 0 | 0 | 0 | 0 |
| 0.125 | 1.56E-06 | 0.000914 | 0 | 0.000301 | 0 | 0.0011 | 6 |
| 0.125 | 5.62E-07 | 0 | 0 | 0.000301 | 18.4 | 0 | 0 |
| 0.125 | 1.56E-06 | 0 | 0.005257 | 0 | 18.4 | 0 | 6 |
| 0 | 1.56E-06 | 0.000914 | 0.005257 | 0.000128 | 0 | 0.000622 | 0 |
| 0.125 | 0 | 0 | 0.005257 | 0 | 8.924 | 0 | 6 |
| 0 | 1.56E-06 | 0 | 0.005257 | 0.000301 | 0 | 0 | 6 |
| 0 | 0 | 0 | 0 | 0 | 0 | 0 | 0 |
| 0.125 | 1.56E-06 | 0.000914 | 0.005257 | 0.000301 | 18.4 | 0.0011 | 6 |
| 0 | 1.56E-06 | 0 | 0 | 0.000301 | 18.4 | 0 | 6 |
| 0.125 | 1.56E-06 | 0.000914 | 0.005257 | 0.000301 | 18.4 | 0.0011 | 0 |
| 0.125 | 0 | 0 | 0.005257 | 0 | 0 | 0 | 0 |
| 0.125 | 0 | 0 | 0 | 0.000301 | 18.4 | 0.0011 | 6 |
| 0 | 0 | 0.000914 | 0 | 0.000301 | 18.4 | 0.0011 | 0 |
| 0 | 0 | 0.000914 | 0 | 0 | 0 | 0 | 0 |
| 0 | 0 | 0 | 0.005257 | 0.000301 | 0 | 0.0011 | 6 |
| 0.125 | 8.19E-07 | 0.000914 | 0 | 0 | 18.4 | 0 | 0 |
| 0 | 0 | 0.000914 | 0.005257 | 0 | 18.4 | 0.0011 | 0 |
| 0.06875 | 1.56E-06 | 0.000914 | 0.005257 | 0.000301 | 18.4 | 0 | 0 |
| 0.125 | 1.56E-06 | 0.000914 | 0 | 0.000301 | 0 | 0.000347 | 6 |
| 0 | 0 | 0.000914 | 0 | 0 | 18.4 | 0.000407 | 6 |
| 0 | 0 | 0.000393 | 0.005257 | 0 | 0 | 0 | 6 |
| 0.006987 | 2.34E-07 | 7.32E-05 | 0.00439 | 0.000125 | 0 | 0.00066 | 6 |
| 0.125 | 0 | 0.000914 | 0.005257 | 0 | 18.4 | 0 | 6 |
| 0.125 | 1.56E-06 | 0 | 0 | 0 | 11.04 | 0 | 6 |
| 0.125 | 1.56E-06 | 0 | 0 | 0.000301 | 0 | 0 | 0 |
| 0.125 | 1.56E-06 | 0.000914 | 0.002497 | 0 | 0 | 0 | 6 |
| 0.125 | 0 | 0.000914 | 0.005257 | 0.000187 | 6.9 | 0 | 3.3 |
| 0.05625 | 0 | 0 | 0 | 0 | 18.4 | 0.00044 | 0 |
| 0.125 | 1.56E-06 | 0 | 0.001866 | 0 | 18.4 | 0.00059 | 0 |
| 0 | 1.56E-06 | 0.000914 | 0 | 0.000135 | 0 | 0.0011 | 6 |
| 0.125 | 0 | 0 | 0 | 0 | 0 | 0.0011 | 0 |
| 0 | 1.26E-06 | 0.000199 | 0.002277 | 6.56E-05 | 18.4 | 0.0011 | 5.88 |
| 0 | 0 | 0.000914 | 0.005257 | 0 | 9.2 | 0 | 0 |
| 0.075 | 1.56E-06 | 0.000914 | 0.002056 | 0.000103 | 0 | 0.0011 | 2.11026 |
| 0 | 1.56E-06 | 0 | 0.005257 | 0.000301 | 18.4 | 0 | 6 |

| | | | | | | | |
|---|---|---|---|---|---|---|---|
| 0 | 1.56E-06 | 0.000914 | 0 | 0 | 18.4 | 0.0011 | 6 |
| 0.059375 | 1.56E-06 | 0.000914 | 0.002707 | 0.000301 | 18.4 | 0.0011 | 3.09 |
| 0.125 | 8.58E-07 | 0.000914 | 0.005257 | 0 | 0 | 0.0011 | 3.15 |
| 0 | 1.56E-06 | 0 | 0 | 0.000301 | 18.4 | 0.0011 | 6 |
| 0.125 | 0 | 0 | 0.005257 | 0.000301 | 18.4 | 0 | 6 |
| 0.053125 | 1.56E-06 | 0.000914 | 0.005257 | 0 | 0 | 0.0011 | 6 |
| 0.125 | 0 | 0.000914 | 0.005257 | 0 | 0 | 0.0011 | 3.48 |
| 0 | 1.56E-06 | 0 | 0 | 0.000301 | 18.4 | 0 | 0 |
| 0 | 0 | 0 | 0 | 0.000301 | 0 | 0 | 0 |
| 0.061875 | 1.56E-06 | 0 | 0.005257 | 0 | 18.4 | 0 | 0 |
| 0 | 1.56E-06 | 0 | 0.005257 | 0 | 0 | 0.00066 | 2.7 |
| 0 | 1.56E-06 | 0.000443 | 0 | 0 | 18.4 | 0.0011 | 0 |
| 0.125 | 0 | 0.000521 | 0 | 0 | 8.648 | 0.0011 | 6 |
| 0.125 | 1.56E-06 | 0 | 0 | 0.000301 | 18.4 | 0 | 6 |
| 0.125 | 6.32E-07 | 0.000914 | 0 | 0.000301 | 0 | 0.0011 | 6 |
| 0 | 0 | 0.000914 | 0 | 0.000301 | 0 | 0 | 6 |
| 0 | 1.56E-06 | 0.000914 | 0.002497 | 0 | 0 | 0 | 6 |
| 0 | 0 | 0.000393 | 0.005257 | 0.000301 | 18.4 | 0.00066 | 6 |
| 0 | 1.56E-06 | 0.000914 | 0.005257 | 0.000301 | 0 | 0.0011 | 0 |
| 0 | 0 | 0 | 0.005257 | 0.000135 | 0 | 0.0011 | 0 |
| 0.125 | 1.56E-06 | 0.000914 | 0.005257 | 0.000301 | 0 | 0 | 0 |
| 0.125 | 0 | 0.000914 | 0 | 0 | 18.4 | 0.0011 | 0 |
| 0 | 0 | 0 | 0 | 0 | 0 | 0 | 0 |
| 0 | 0 | 0 | 0 | 0 | 0 | 0 | 0 |
| 0.125 | 1.56E-06 | 0 | 0.005257 | 0.000301 | 0 | 0.0011 | 6 |
| 0.125 | 0 | 0.000914 | 0.005257 | 0.000301 | 18.4 | 0 | 0 |
| 0.125 | 1.56E-06 | 0 | 0 | 0 | 0 | 0 | 0 |
| 0.125 | 1.56E-06 | 0 | 0 | 0 | 0 | 0.0011 | 0 |
| 0 | 4.99E-07 | 0.000914 | 0.005257 | 0.000301 | 0 | 0 | 0 |
| 0.125 | 1.56E-06 | 0.000457 | 0.005257 | 0 | 0 | 0 | 0 |
| 0.04375 | 7.2E-07 | 0 | 0.003128 | 0 | 18.4 | 0.0011 | 6 |
| 0 | 0 | 0.000914 | 0.005257 | 0 | 0 | 0 | 6 |
| 0.125 | 1.56E-06 | 0.000914 | 0 | 0 | 0 | 0 | 6 |
| 0 | 0 | 0 | 0.005257 | 0.000194 | 18.4 | 0 | 0 |
| 0 | 1.56E-06 | 0.000914 | 0 | 0.000301 | 0 | 0 | 0 |
| 0.125 | 1.56E-06 | 0.000914 | 0 | 0.000301 | 8.74 | 0 | 6 |
| 0.125 | 1.56E-06 | 0 | 0.00276 | 0.000301 | 0 | 0.0011 | 6 |
| 0.125 | 1.56E-06 | 0.000914 | 0 | 0.000301 | 18.4 | 0.0011 | 6 |
| 0 | 8.11E-07 | 0 | 0 | 0 | 10.58 | 0 | 0 |
| 0.04375 | 0 | 0 | 0 | 0.000301 | 18.4 | 0 | 6 |
| 0 | 0 | 0.000914 | 0.005257 | 0 | 0 | 0.0011 | 0 |
| 0 | 1.56E-06 | 0.000914 | 0.005257 | 0.000301 | 12.88 | 0.000627 | 6 |
| 0.125 | 0 | 0 | 0.005257 | 0.000301 | 0 | 0.000622 | 0 |
| 0 | 1.56E-06 | 0 | 0 | 0 | 0 | 0 | 0 |
| 0 | 0 | 0 | 0 | 0 | 0 | 0 | 0 |
| 0 | 1.56E-06 | 0.000503 | 0.005257 | 0.000301 | 0 | 0.0011 | 6 |
| 0.125 | 0 | 0 | 0.005257 | 0 | 18.4 | 0.0011 | 0 |

| | | | | | | | |
|---|---|---|---|---|---|---|---|
| 0.125 | 1.56E-06 | 0.000914 | 0.005257 | 0.000301 | 18.4 | 0 | 6 |
| 0.125 | 1.56E-06 | 0.000914 | 0.001709 | 0.000301 | 0 | 0.0011 | 0 |
| 0.06625 | 1.56E-06 | 0.000914 | 0.005257 | 0.000301 | 0 | 0 | 0 |
| 0.125 | 0 | 0.000544 | 0 | 0.000301 | 18.4 | 0.0011 | 2.7 |
| 0.0625 | 1.56E-06 | 0 | 0 | 0.000114 | 7.36 | 0 | 6 |
| 0 | 1.56E-06 | 0 | 0.005257 | 0.000301 | 18.4 | 0.0011 | 2.64 |
| 0.125 | 0 | 0.000914 | 0.005257 | 0 | 18.4 | 0.000589 | 6 |
| 0.125 | 0 | 0.000914 | 0 | 0.000157 | 0 | 0 | 0 |
| 0.125 | 0 | 0 | 0.005257 | 0.000301 | 0 | 0 | 6 |
| 0 | 0 | 0.000914 | 0.005257 | 0 | 18.4 | 0.0011 | 6 |
| 0.125 | 1.56E-06 | 0 | 0.005257 | 0 | 6.44 | 0.0011 | 0 |
| 0.125 | 7.02E-07 | 0.000914 | 0.002523 | 0.000135 | 18.4 | 0 | 6 |
| 0.125 | 0 | 0 | 0.005257 | 0 | 0 | 0.0011 | 6 |
| 0 | 0 | 0.000914 | 0.005257 | 0.000301 | 0 | 0.0011 | 0 |
| 0 | 0 | 0.000914 | 0 | 0 | 0 | 0.0011 | 6 |
| 0 | 1.56E-06 | 0.000914 | 0 | 0.000301 | 9.2 | 0 | 0 |
| 0 | 0 | 0 | 0.005257 | 0.000301 | 0 | 0.0011 | 6 |
| 0.125 | 1.56E-06 | 0 | 0.005257 | 0.000301 | 18.4 | 0 | 0 |
| 0.07 | 0 | 0.000521 | 0 | 0.000301 | 0 | 0 | 6 |
| 0 | 0 | 0.000914 | 0 | 0.000301 | 18.4 | 0.0011 | 2.88 |
| 0.125 | 1.56E-06 | 0 | 0 | 0 | 18.4 | 0 | 0 |
| 0 | 0 | 0.000914 | 0 | 0.000301 | 18.4 | 0 | 0 |
| 0 | 1.56E-06 | 0 | 0 | 0 | 18.4 | 0.0011 | 0 |
| 0.125 | 1.56E-06 | 0.000297 | 0 | 0.0002 | 0 | 0.0011 | 0 |
| 0 | 0 | 0 | 0 | 0 | 0 | 0 | 0 |
| 0 | 0 | 0 | 0 | 0 | 18.4 | 0 | 6 |
| 0.125 | 0 | 0 | 0.005257 | 0.000301 | 18.4 | 0 | 0 |
| 0 | 0 | 0 | 0 | 0 | 0 | 0 | 0 |
| 0.125 | 6.08E-07 | 0 | 0.005257 | 0.000196 | 18.4 | 0.0011 | 6 |
| 0.125 | 1.56E-06 | 0.000914 | 0.005257 | 0 | 0 | 0 | 0 |
| 0 | 7.8E-07 | 0.000498 | 0 | 0 | 0 | 0.000495 | 3.9 |
| 0.125 | 1.56E-06 | 0.000914 | 0.002629 | 0 | 8.648 | 0.0011 | 0 |
| 0.025923 | 1.12E-06 | 0.000914 | 0.00481 | 0 | 18.4 | 0.000163 | 5.85 |
| 0 | 1.56E-06 | 0 | 0 | 0 | 18.4 | 0.0011 | 6 |
| 0.125 | 1.56E-06 | 0.000914 | 0 | 0 | 0 | 0.0011 | 6 |
| 0 | 1.56E-06 | 0 | 0.005257 | 0 | 18.4 | 0.00055 | 6 |
| 0 | 1.56E-06 | 0.000914 | 0 | 0.000301 | 18.4 | 0 | 0 |
| 0 | 0 | 0 | 0 | 0.000301 | 0 | 0 | 0 |
| 0.05625 | 0 | 0.000914 | 0 | 0 | 18.4 | 0.0011 | 0 |
| 0.125 | 0 | 0 | 0 | 0.000301 | 0 | 0.0011 | 0 |
| 0.125 | 0 | 0 | 0 | 0 | 0 | 0 | 0 |
| 0 | 1.56E-06 | 0.000914 | 0 | 0.000301 | 18.4 | 0.0011 | 0 |
| 0 | 1.56E-06 | 0.000914 | 0 | 0.000301 | 0 | 0.0011 | 0 |
| 0 | 1.56E-06 | 0.000914 | 0.005257 | 0 | 18.4 | 0.0011 | 6 |
| 0.09625 | 7.8E-08 | 0.000914 | 0.000394 | 0.000167 | 13.8 | 0 | 0 |
| 0 | 1.56E-06 | 0.000914 | 0 | 0 | 18.4 | 0 | 0 |
| 0.125 | 0 | 0 | 0.005257 | 0.000301 | 18.4 | 0.0011 | 6 |

| | | | | | | | |
|---|---|---|---|---|---|---|---|
| 0.125 | 1.56E-06 | 0 | 0 | 0.000301 | 18.4 | 0 | 3.15 |
| 0.125 | 0 | 0 | 0 | 0 | 0 | 0 | 6 |
| 0 | 0 | 0 | 0 | 0 | 0 | 0 | 0 |
| 0.125 | 0 | 0 | 0 | 0 | 18.4 | 0 | 6 |
| 0.125 | 1.56E-06 | 0 | 0.005257 | 0.000301 | 18.4 | 0.0011 | 0 |
| 0.125 | 1.56E-06 | 0.000914 | 0 | 0 | 18.4 | 0 | 6 |
| 0 | 0 | 0 | 0 | 0 | 0 | 0 | 0 |
| 0 | 8.74E-07 | 0.000448 | 0.005257 | 0.000301 | 18.4 | 0 | 6 |
| 0.125 | 1.56E-06 | 0.000599 | 0.003458 | 0.000301 | 1.84 | 0.000724 | 2.20556 |
| 0.06875 | 0 | 0 | 0.005257 | 0.000301 | 9.2 | 0 | 3.72 |
| 0 | 0 | 0 | 0 | 0.000301 | 18.4 | 0.000633 | 0 |
| 0.125 | 0 | 0.000914 | 0 | 0 | 0 | 0 | 0 |
| 0.125 | 0 | 0 | 0 | 0.000135 | 18.4 | 0.0011 | 3.51 |
| 0.125 | 1.56E-06 | 0.000539 | 0 | 5.02E-05 | 10.12 | 0 | 0 |
| 0 | 0 | 0 | 0 | 0 | 0 | 0 | 0 |
| 0.125 | 1.56E-06 | 0.000914 | 0.005257 | 0.000301 | 0 | 0 | 0 |
| 0 | 0 | 0 | 0 | 0.000301 | 0 | 0.0011 | 6 |
| 0 | 1.56E-06 | 0.000914 | 0.005257 | 0 | 0 | 0 | 6 |
| 0 | 1.56E-06 | 0.000914 | 0 | 0 | 0 | 0.00077 | 0 |
| 0 | 0 | 0 | 0 | 0 | 18.4 | 0.0011 | 0 |
| 0 | 1.56E-06 | 0 | 0.005257 | 0.000301 | 0 | 0.0011 | 0 |
| 0.125 | 0 | 0.000516 | 0.005257 | 0.000301 | 18.4 | 0.0011 | 0 |
| 0.125 | 1.56E-06 | 0.000914 | 0 | 0 | 18.4 | 0.0011 | 6 |
| 0 | 1.56E-06 | 0.000914 | 0.005257 | 0.000301 | 7.82 | 0 | 6 |
| 0 | 1.56E-06 | 0 | 0.005257 | 0 | 18.4 | 0.0011 | 6 |
| 0 | 0 | 0.000914 | 0.002891 | 0.000301 | 18.4 | 0 | 0 |
| 0 | 0 | 0 | 0 | 0 | 0 | 0.0011 | 0 |
| 0.125 | 1.56E-06 | 0.000914 | 0.005257 | 0.000301 | 18.4 | 0 | 6 |
| 0.125 | 0 | 0.000914 | 0 | 0 | 0 | 0.0011 | 0 |
| 0 | 1.56E-06 | 0 | 0.005257 | 0 | 0 | 0 | 0 |
| 0.09625 | 1.01E-06 | 0.000506 | 0.001072 | 0.000301 | 18.4 | 0.0011 | 2.81229 |
| 0.125 | 0 | 0.000914 | 0.005257 | 0.000301 | 0 | 0.0011 | 6 |
| 0.06125 | 1.56E-06 | 0 | 0 | 0.000301 | 0 | 0.0011 | 0 |
| 0 | 0 | 0 | 0 | 0.000301 | 0 | 0.000585 | 6 |
| 0 | 0 | 0.000489 | 0.002629 | 0.000301 | 0 | 0 | 0 |
| 0.125 | 0 | 0 | 0.005257 | 0 | 18.4 | 0 | 0 |
| 0.125 | 0 | 0 | 0 | 0 | 18.4 | 0.0011 | 6 |
| 0 | 0 | 0.000914 | 0 | 0 | 18.4 | 0.0011 | 6 |
| 0 | 1.56E-06 | 0 | 0.005257 | 0 | 18.4 | 0 | 6 |
| 0 | 0 | 0 | 0.002392 | 0.000301 | 0 | 0.0011 | 6 |
| 0 | 0 | 0.000914 | 0 | 0.000301 | 0 | 0.0011 | 6 |
| 0 | 1.56E-06 | 0 | 0.005257 | 0 | 0 | 0.0011 | 0 |
| 0 | 0 | 0.000914 | 0.005257 | 0.000301 | 18.4 | 0 | 6 |
| 0.125 | 1.56E-06 | 0 | 0 | 0.000301 | 0 | 0.0011 | 6 |
| 0 | 9.05E-07 | 0 | 0.005257 | 0 | 0 | 0 | 0 |
| 0 | 9.7E-07 | 0.000914 | 0.005257 | 0 | 18.4 | 0.0011 | 0 |
| 0 | 0 | 0 | 0.005257 | 0 | 0 | 0 | 6 |

| | | | | | | | |
|---|---|---|---|---|---|---|---|
| 0.125 | 0 | 0 | 0.002891 | 0 | 18.4 | 0 | 0 |
| 0 | 1.56E-06 | 0 | 0.005257 | 0.000114 | 0 | 0 | 6 |
| 0.125 | 0 | 0.000914 | 0.005257 | 0 | 0 | 0.0011 | 6 |
| 0.125 | 0 | 0.000914 | 0.005257 | 0.000301 | 18.4 | 0.0011 | 0 |
| 0.065625 | 0 | 0.000914 | 0.005257 | 0.000164 | 0 | 0.0011 | 6 |
| 0.059439 | 1.09E-06 | 0.000914 | 0 | 0.000181 | 18.4 | 0 | 6 |
| 0.125 | 1.56E-06 | 0.000914 | 0.005257 | 0 | 0 | 0.0011 | 6 |
| 0.125 | 0 | 0.000914 | 0.005257 | 0 | 0 | 0.000572 | 0 |
