## Supplemental code 1 for "Micronutrient Optimization Using Design of Experiments Approach in Tissue Engineered Articular Cartilage for Production of Type II Collagen"

```
from opentrons import protocol_api
#sample collection and feeding: 300ul tips, changed api level to 2.6 from
9-9 protocol
# metadata
metadata = {
    'protocolName': 'Sample_and_Feed',
    'author': 'Name ',
    'description': 'sample media and replace with fresh media',
    'apiLevel': '2.11'
}


def run(protocol: protocol_api.ProtocolContext):
#Labware Definitions
    cellplate1 =
protocol.load_labware('greinerbioone_96_wellplate_280ul','10')
    whiteplate1 = protocol.load_labware('corning_96_wellplate_360ul_flat',
'8')
    stockplate1 = protocol.load_labware('vwr_96_wellplate_2500ul', '11')
    cellplate2 =
protocol.load_labware('greinerbioone_96_wellplate_280ul','4')
    whiteplate2 = protocol.load_labware('corning_96_wellplate_360ul_flat',
'2')
    stockplate2 = protocol.load_labware('vwr_96_wellplate_2500ul', '5')
    waste2 = protocol.load_labware('corning_96_wellplate_360ul_flat', '1')
    waste1 = protocol.load_labware('corning_96_wellplate_360ul_flat', '7')
    tiprack1 = protocol.load_labware('opentrons_96_tiprack_300ul', '9')
    tiprack2 = protocol.load_labware('opentrons_96_tiprack_300ul', '6')


# pipettes and tipracks
    right_pipette = protocol.load_instrument(
         'p300_multi_gen2', mount='right', tip_racks=[tiprack1, tiprack2])
#plate columns in use
    plate_loc = ['A2', 'A3', 'A4', 'A5', 'A6', 'A7', 'A8',
'A9','A10','A11']

#functions
    for i in range (20):
            if i < 10:
                right_pipette.pick_up_tip()
                right_pipette.well_bottom_clearance.aspirate = 2
                right_pipette.flow_rate.aspirate = 40
                right_pipette.aspirate(150,
cellplate1.wells_by_name()[plate_loc[i]])
                right_pipette.dispense(130,
waste1.wells_by_name()[plate_loc[i]])
                right_pipette.dispense(20,
whiteplate1.wells_by_name()[plate_loc[i]])
                right_pipette.touch_tip(v_offset=-2)
                right_pipette.well_bottom_clearance.aspirate = 2
                right_pipette.aspirate(150,
stockplate1.wells_by_name()[plate_loc[i]])
                right_pipette.dispense(150,
cellplate1.wells_by_name()[plate_loc[i]])
                right_pipette.drop_tip()
```

```
            if 20 > i >=10:
                right_pipette.pick_up_tip()
                right_pipette.well_bottom_clearance.aspirate = 2
                right_pipette.flow_rate.aspirate = 40
                right_pipette.aspirate(150,
cellplate2.wells_by_name()[plate_loc[i-10]])
                right_pipette.dispense(130,
waste2.wells_by_name()[plate_loc[i-10]])
                right_pipette.dispense(20,
whiteplate2.wells_by_name()[plate_loc[i-10]])
                right_pipette.touch_tip(v_offset=-2)
                right_pipette.well_bottom_clearance.aspirate = 2
                right_pipette.aspirate(150,
stockplate2.wells_by_name()[plate_loc[i-10]])
                right_pipette.dispense(150,
cellplate2.wells_by_name()[plate_loc[i-10]])
                right_pipette.drop_tip(
```
