## Supplementary material for "Micronutrient Optimization Using Design of Experiments Approach in Tissue Engineered Articular Cartilage for Production of Type II Collagen": other supplemental

| Term | Conditions (µg/ml) | | | | |
|---|---|---|---|---|---|
| | **12** | **25** | **52** | **72** | **89** |
| **Lin** | 6.6E-09 | 3.3E-08 | 1.1E-08 | 6.8E-09 | 5.6E-09 |
| **Cr** | 3.9E-04 | 6.8E-04 | 4.9E-05 | 6.6E-04 | 1.1E-03 |
| **Co** | 5.8E-06 | 1.2E-04 | 1.3E-06 | 1.8E-06 | 5.5E-06 |
| **Cu** | 2.1E-01 | 2.2E-08 | 6.7E-01 | 4.6E-02 | 5.0E-02 |
| **I** | 9.0E-02 | 1.5E-02 | 9.2E-02 | 9.2E-02 | 9.0E-02 |
| **Mn** | 1.2E-02 | 4.5E-05 | 4.7E-04 | 8.6E-03 | 3.8E-03 |
| **Mo** | 1.9E-03 | 1.5E-03 | 2.0E-03 | 2.0E-03 | 1.1E-03 |
| **Thy** | 2.5E-02 | 2.6E-02 | 2.4E-02 | 2.4E-02 | 2.4E-02 |
| **Vit A** | 8.4E-11 | 3.0E-11 | 2.5E-11 | 1.0E-10 | 9.8E-11 |
| **Vit B12** | 9.1E-07 | 6.3E-12 | 1.3E-06 | 9.1E-07 | 9.7E-07 |
| **Vit B7** | 2.8E-03 | 2.3E-03 | 3.0E-03 | 9.3E-05 | 1.9E-03 |
| **Vit D** | 7.2E-06 | 9.9E-10 | 1.7E-05 | 8.7E-08 | 8.7E-08 |
| **Vit E** | 1.5E-08 | 5.9E+01 | 1.6E-08 | 1.5E-08 | 2.0E-08 |
| **Vit K** | 4.7E-06 | 5.0E-12 | 5.6E-06 | 2.8E-05 | 9.0E-06 |
| **Zn** | 3.6E-03 | 2.3E+00 | 1.8E-03 | 4.4E-03 | 1.8E-03 |
| ***Desirability*** | *0.835* | *0.595* | *0.831* | *0.829* | *0.827* |

**Supplemental Table 2**: Optimal predicted combinations of vitamins and minerals as derived by DoE.

a.

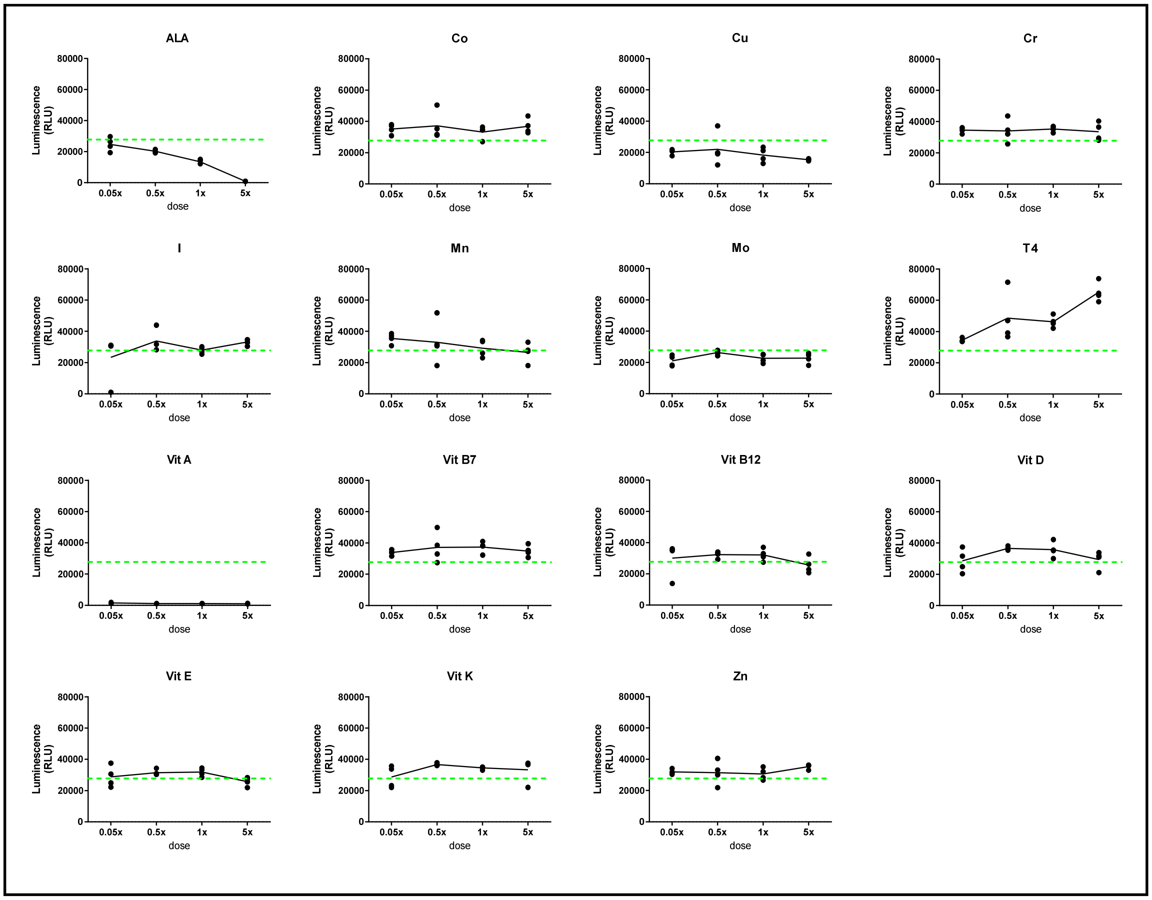


b.

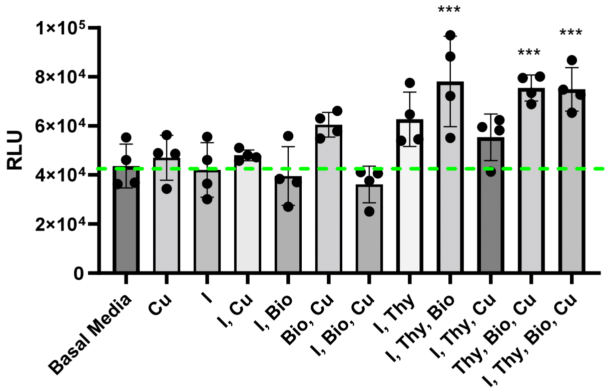


**Supplemental Fig 1: a, b** Primary COL2A1-GLuc rabbit chondrocytes were seeded in aggregate culture in basal chondrogenic media supplemented with different concentrations of a single vitamin or mineral **(a)** or combinations **(b)**. Media was assessed for luminescence and results are shown for day 21. Individual values for 4 replicates are shown with green dashed line indicating basal media mean.

Error bars **(b)** indicate standard deviation and *** indicate $p < 0.001$ vs. basal media control.

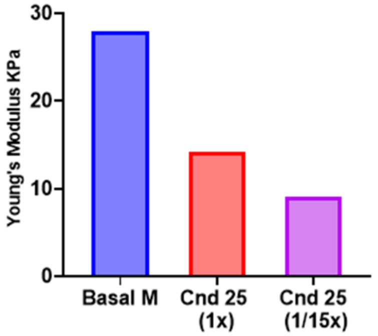


**Supplemental Fig 2:** COL2A1-GLuc primary rabbit chondrocytes were cultured in custom in house bioreactors. At day 22, biopsy punches of engineered sheet were assessed via compression testing  and young's modulus calculated.
